# Control of CDK activity and the cell cycle by CKS proteins

**DOI:** 10.1101/2025.07.04.663185

**Authors:** Joseph F. Curran, Souradeep Basu, Tania Auchynnikava, Paul Nurse

## Abstract

The eukaryotic cell cycle is orchestrated by the activity of Cyclin-Dependent Kinases (CDKs)^1^. Although most mechanisms controlling CDK activity are well understood, one class of proteins - the CDK regulatory subunit (CKS) proteins - remain highly enigmatic. There is no generally accepted consistent molecular or functional characterisation of CKS *in vivo*, despite being essential across eukaryotes and implicated in a range of cell cycle processes and cancer^2,3^. Here, we provide a unifying framework for CKS function. We show in a single, genetically tractable system, the fission yeast, that CKS regulates the onset and progression of both S-phase and mitosis. We find that CKS modulates the phosphorylation of more than 200 CDK phosphosites *in vivo*, located on 133 substrate proteins. These are involved in diverse processes across the entire cell cycle, including DNA replication, chromosome condensation, the spindle assembly checkpoint, and the metaphase-anaphase transition. We demonstrate that CKS enhances CDK activity to drive phosphorylation of sites with low affinity for CDK. This acts both on specific substrates, likely by stabilizing CDK-substrate interactions, and through control of overall CDK activity, by regulating its interaction with Wee1 and Cdc25. Our findings establish CKS proteins as major multifaceted regulators of the cell cycle, operating through the global and local control of CDK activity.

## Introduction

Cyclin-Dependent Kinases (CDKs) phosphorylate hundreds of proteins to bring about the events of the eukaryotic cell cycle ^4,5^. CDK activity is dependent on the formation of Cyclin-CDK complexes and is modulated by inhibitory CDK-Y15 phosphorylation^6,7^. CDKs also bind a <u>C</u>D<u>K</u> regulatory <u>S</u>ubunit (CKS), forming tripartite Cyclin-CDK-CKS complexes^8–11^. CKS proteins are small (9-13 kDa), essential, and highly conserved, yet their contribution to CDK regulation remains uncertain. CKS was discovered as *suc1* in the fission yeast *Schizosaccharomyces pombe*^12,13^. Subsequently, homologues were identified in *Saccharomyces cerevisiae,* where it was named *CKS*, and then in metazoans, with two isoforms - CKS1B and CKS2 in humans^9–11,14^. Despite their sequence conservation and ubiquitous association with CDK, investigations in different experimental systems have reported highly diverse effects upon loss of function, including defects or cell cycle arrests in G1/S^15–18^, G2^9,13,15,17,19,20^ and mitosis^9,19,21–24^. Both increased and decreased CKS protein levels impact cell cycle control^11,13,20,25,26^, and elevated CKS expression is associated with tumour aggressiveness and malignant progression of multiple cancers^3,27,28^. As a result of this complexity, it is unclear if CKS performs different roles in particular organisms and cell types, and there is no generally accepted view of how CKS proteins impact cell cycle control.

There have also been proposals of many different molecular mechanisms of CKS function. These include roles as structural adaptors promoting transcriptional regulation^17,29^ or degradation of proteins such as the CDK inhibitor p27^Kip1^ by E2F^16,30,31^ and cyclins by the APC/C^23,32–34^. *In vitro* studies using reporter substrates have shown that CKS proteins can enhance multisite phosphorylation cascades by docking to CDK substrates via a conserved phospho-threonine binding pocket^22,35–38^. This has led to the suggestion that CKS proteins act as accessory proteins which enhance multisite phosphorylation of CDK substrates^39–41^. However, the impacts of CKS proteins on global CDK substrate phosphorylation *in vivo*, on CDK regulation and on cell cycle control remain unknown. To address these questions, we have undertaken a systematic characterisation of the biological and molecular functions of CKS proteins *in vivo* in fission yeast and propose a unifying framework for the role of CKS proteins in control of the eukaryotic cell cycle.

## Results

### The essential functions of CKS proteins are conserved

Given the diversity of functions reported for CKS proteins, we tested if their essential functions are conserved between fission yeast and other eukaryotes. One copy of *suc1* in diploid fission yeast cells was replaced with genes encoding either the human *Hs*CKS1B or *Hs*CKS2 homologues or the budding yeast *Sc*CKS1 homologue. Sporulation of these heterozygotes revealed that any one of these CKS proteins could rescue deletion of *suc1*Δ resulting in similar growth rates and cell division sizes to wildtype cells (Fig. 1a,b, Extended Data Fig. 1a,b). *CKS1Δ* in *S. cerevisiae* can also be rescued by *suc1*, HsCKS1B or HsCKS2^11^. Therefore, the essential functions of Suc1 are conserved between fission yeast, budding yeast and human CKS proteins.

**Fig. 1:**
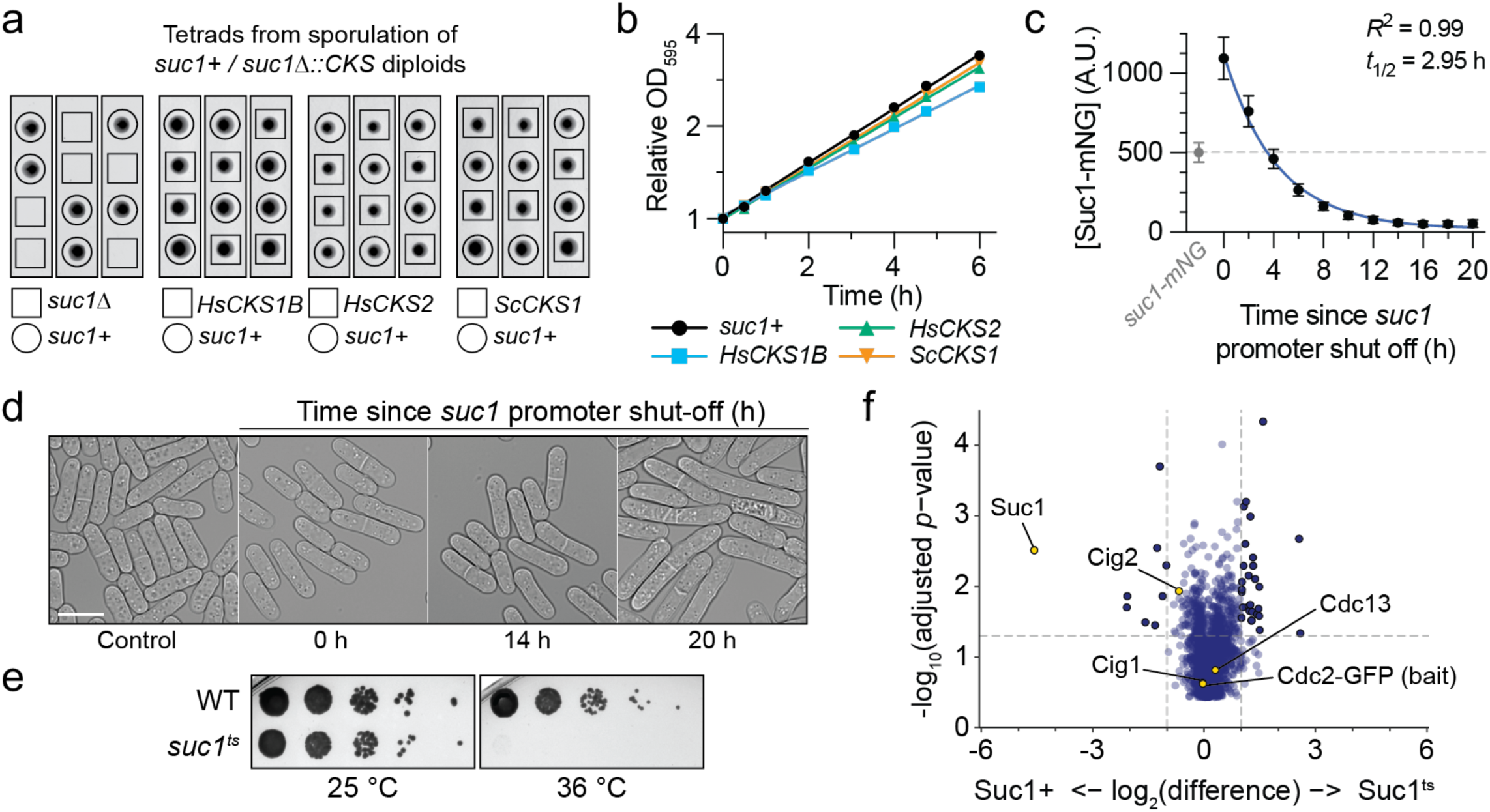
*suc1^ts^* is a lethal temperature-sensitive allele of the conserved CKS protein Suc1. **a**, Representative tetrads of haploids dissected from sporulation of heterozygous diploids. *suc1* allele is indicated by circles/boxes, as determined by replica plating and a selectable marker. **b**, Increase in OD_595_ over time relative to time 0. Mean and range shown for *n* = 2 biological repeats. Error bars are masked by the point markers. **c**, Whole cell concentration of Suc1-mNG over time after repression of *suc1* promoter. A.U., arbitrary units. Mean and s.d. shown for *n* > 1000 cells per timepoint; representative of *n* = 2 biological repeats. Least-squares regression fit of a one-phase exponential decay. **d**, Example images of cells following repression of *suc1* expression. Scale bar = 10 µm. **e**, Representative serial dilution assay for colony formation. **f**, Multiple testing corrected T-test significance (Q values) vs fold change in protein abundance detected by mass spectrometry following Cdc2-GFP immunoprecipitation. *n* = 3 biological replicates per strain.

To conditionally inactivate Suc1, we placed *suc1* under the control of a thiamine-repressible promoter. De-repressed expression resulted in two-fold overexpression relative to endogenous levels and delayed cell cycle progression as determined by increased length at cell division (Fig. 1c,d, Extended Data Fig. 1c). Promoter repression resulted in a decrease in Suc1 concentration over time, and this under-expression initially advanced cells into mitosis, consistent with dose-dependent modulation of cell cycle control^13,20,26^. However, cells continued cycling for a further 4 generations (12 h) before defects in cell cycle progression and viability became apparent, despite Suc1 levels falling to 10% of endogenous levels (Fig. 1d, Extended Data Fig. 1c,d). We conclude that only a low level of Suc1 is required for cell viability, and that residual CKS levels from experimental approaches such as repressible promoters and siRNA knockdowns may still be sufficient for CKS function that is biologically significant.

To rapidly inactivate Suc1, we performed a targeted mutagenic screen based on an *S. cerevisiae CKS1-ts38* allele^15^ (Methods) and isolated a temperature-sensitive mutant which was inviable at 36 °C, *suc1(A32T, E35Y, M42T, N57D, L74H)*, hereafter *suc1^ts^* (Fig. 1e). Three mutated residues in *suc1^ts^* lie close to the CDK-Suc1 binding interface, so we tested if Suc1 binding to CDK was compromised at the restrictive temperature. Immunoprecipitation of Cdc2 (*S. pombe* CDK1) followed by mass spectrometry revealed a >20-fold reduction in associated Suc1 in the *suc1^ts^* mutant (Fig. 1f, Supplementary Table 1). There were no changes in the association of Cdc2 with the cyclins, Cig1, Cig2 and Cdc13. Temperature shift of the *suc1^ts^* allele therefore disrupts Suc1 association with Cyclin-CDK, facilitating study of Suc1 function in Cyclin-CDK complexes *in vivo*.

### Suc1 enhances phosphorylation of a subset of CDK phosphosites *in vivo*

We investigated the impact of Suc1 on CDK substrate phosphorylation at the G1/S and G2/M transitions using phosphoproteomics. Cells were synchronised in G1 using nitrogen starvation, or in G2 using inhibition of an ATP analogue-sensitive *cdc2^as^* allele, before temperature shift and release from these blocks. We detected 11,841 and 30,169 phosphosites in the G1/S and G2/M phosphoproteomic experiments respectively. This identified 45 phosphosites at G1/S and 351 at G2/M that we had previously determined to be CDK substrates (ref. 4, and unpublished data), and which increased in phosphorylation by at least 2-fold at these transitions in the *suc1^+^* strain (Extended Data Fig. 2a-d, Methods). The protein abundance of these CDK substrates remained essentially unchanged in both the *suc1^+^* and *suc1^ts^* strains (Extended Data Fig. 2e,g) and phosphorylation intensities at the start of each time course were comparable between strains (Extended Data Fig. 2f,h). Therefore, phosphorylation intensity was normalised to the maximum phosphorylation level in *suc1^+^* cells, to generate a scaled ratio (Methods).

Comparison with the s*uc1^ts^* strain revealed that a substantial subset of CDK sites were hypophosphorylated in the absence of Suc1 function. At G1/S, the phosphorylation of 16 % of sites (7 sites) were strongly inhibited in the *suc1^ts^* strain, and another 20 % (9 sites) were partially inhibited (Fig. 2a,c,e, Supplementary Table 2). At G2/M, 18 % (63 sites) were strongly inhibited and another 41 % (143 sites) were partially inhibited (Fig. 2b,d,f, Supplementary Table 3). In total, 221 CDK phosphosites (out of 390 non-redundant sites, 57 %) were Suc1-dependent, with 70 (18 %) being strongly Suc1-dependent. A further 12 sites at G1/S (27 %) were phosphorylated to greater than 1.5x wildtype levels, suggesting that the presence of Suc1 inhibits phosphorylation of these sites (Fig. 2a,c,e). We conclude that around half of the CDK phosphosites detected *in vivo* are Suc1-dependent for full phosphorylation with about one fifth being highly dependent on Suc1.

**Fig. 2:**
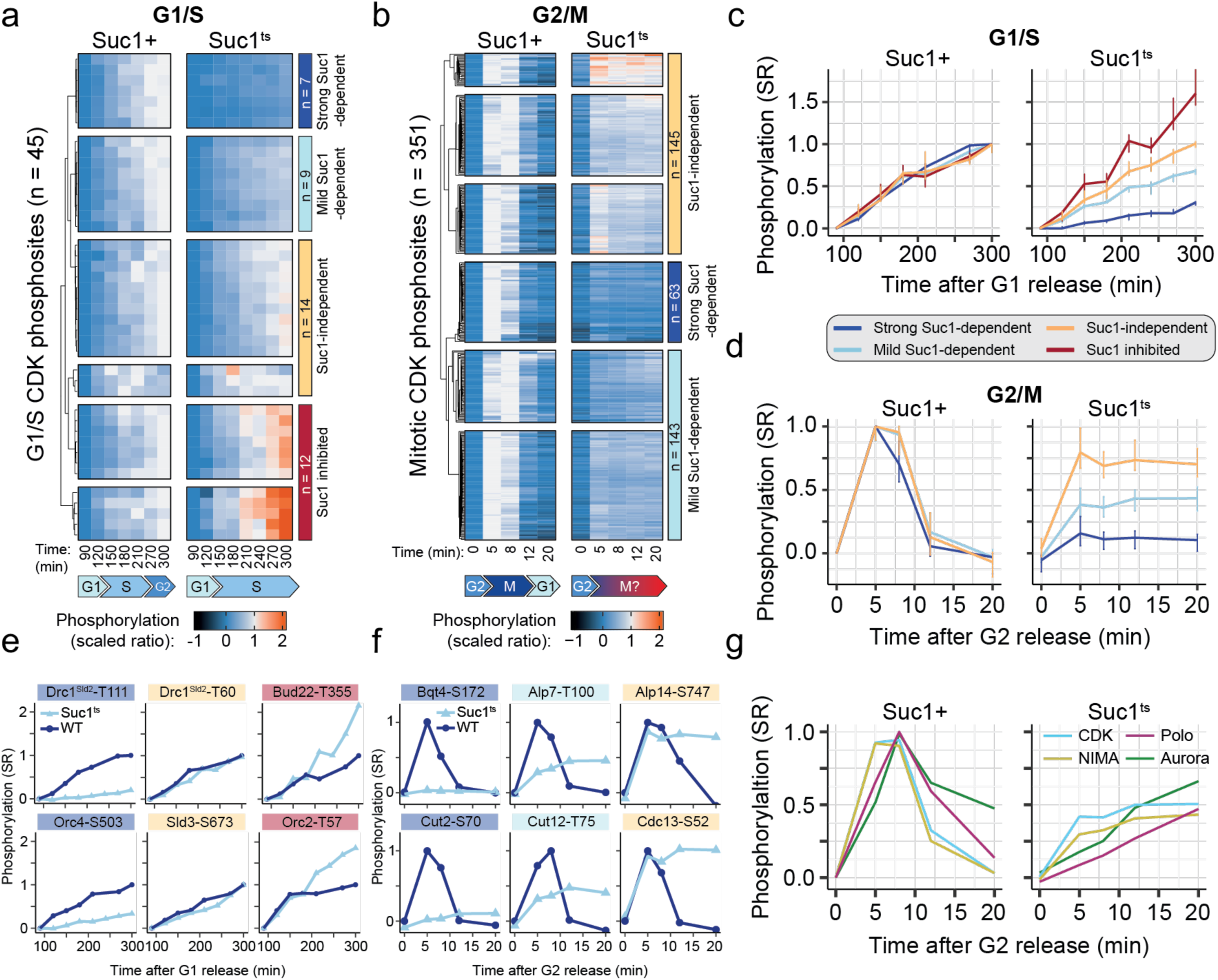
Suc1 modulates phosphorylation of a subset of CDK substrates. Heatmap of the phosphorylation intensity of 45 G1/S (**a**) and 351 G2/M (**b**) CDK phosphosites during progression through S-phase, and mitosis, respectively. Sites were clustered by Euclidean distance, revealing different classes of behaviour. **c** and **d**, Average phosphorylation behaviour (median and IQR) of the classes identified in **a** and **b**, respectively. SR = scaled ratio. **e** and **f**, Representative phosphosites from the major behaviour classes in **a** and **b**, respectively. **g**, Median phosphorylation intensity of putative substrates of CDK (Cdc2; [ST]-P, *n* = 224) Polo (Plo1; [DENQ]-X-[ST]-φ or [ST]-F, *n* = 65), NIMA (Fin1; [LF]-X-X-[ST], *n* = 90) and Aurora (Ark1; R-X-[ST], *n* = 28) kinases during mitosis, based on their consensus motifs^51,85–90^.

### Suc1-dependent substrates are involved in diverse cell cycle processes

The 221 Suc1-dependent phosphosites were found on 133 proteins, 53 of which contained highly Suc1-dependent phosphosites. Of these 133 proteins, 36 are annotated with the Gene Ontology (GO) biological process term “Cell Cycle”, and 29 result in cell cycle arrest when genetically deleted in *S. pombe*^42^. In combination these amount to 51 Suc1-dependent substrates that are directly implicated in progress through the cell cycle. Suc1-dependent substrates were associated with specific cell cycle GO annotations, including centromere localisation, spindle attachment and elongation, checkpoint signalling, the G2/M transition and DNA replication initiation (Extended Data Fig. 2i). We also identified individual Suc1-dependent substrates with well-defined cell cycle roles, including the DNA replication origin firing factor Drc1^Sld2^, the mitotic CDK activating phosphatase Cdc25, the condensin subunit Cut3^Smc4^ and Cut2^Securin^ required for chromosome condensation and sister chromatid separation respectively, and proteins involved in centrosome and kinetochore attachment and dynamics, including Bqt4, Plo1^Polo^ and Pic1^INCENP^, Nsk1, Clp1^Cdc14^ and Alp7 (Fig. 2e,f). Many of the diverse roles reported for CKS proteins can therefore be explained by their wide-ranging effects on the phosphorylation of CDK substrates involved in control and progression through the cell cycle.

CDK activates downstream mitotic kinases to drive the events of mitosis^43–46^, so we also analysed the phosphorylation of putative substrates of the NIMA-related (Fin1), Polo-like (Plo1) and Aurora B (Ark1) kinases. We observed impaired phosphorylation of substrates conforming to the consensus motifs of these kinases, with the strongest effect on Polo kinase substrates (Fig. 2g). This indicates that Suc1 enhances both direct CDK substrate phosphorylation, and the phosphorylation of downstream substrates by other mitotic kinases.

### Suc1 is required for proper execution of S-phase and mitosis

Given the wide influence Suc1 has on CDK substrate phosphorylation we investigated the effects of Suc1 loss of function on progression through the cell cycle. Temperature-shift of asynchronous *suc1^ts^* cells resulted in a range of cell cycle defects (Extended Data Fig. 3a), so we assessed the impact of *suc1^ts^* on the G1/S and G2/M transitions individually. The G1/S transition and S-phase progression was monitored by DNA content analysis following release from G1 arrest via nitrogen starvation. In the *suc1^ts^* mutant, there was a 55-60 min delay in the time taken to complete S-phase (Fig. 3a,b), resulting from a 25 min delay in the time taken for cells to enter S-phase, and a 30 min increase in the average duration of S-phase (Fig. 3c,d, Extended Data Fig. 3b). These defects could result from inefficient origin firing given that Drc1^Sld2^ phosphorylation is dependent on Suc1 (Fig. 2e), and Drc1^Sld2^ phosphorylation is required for initiation of DNA replication^47^. We conclude that Suc1 affects both onset and progression through S-phase.

**Fig. 3:**
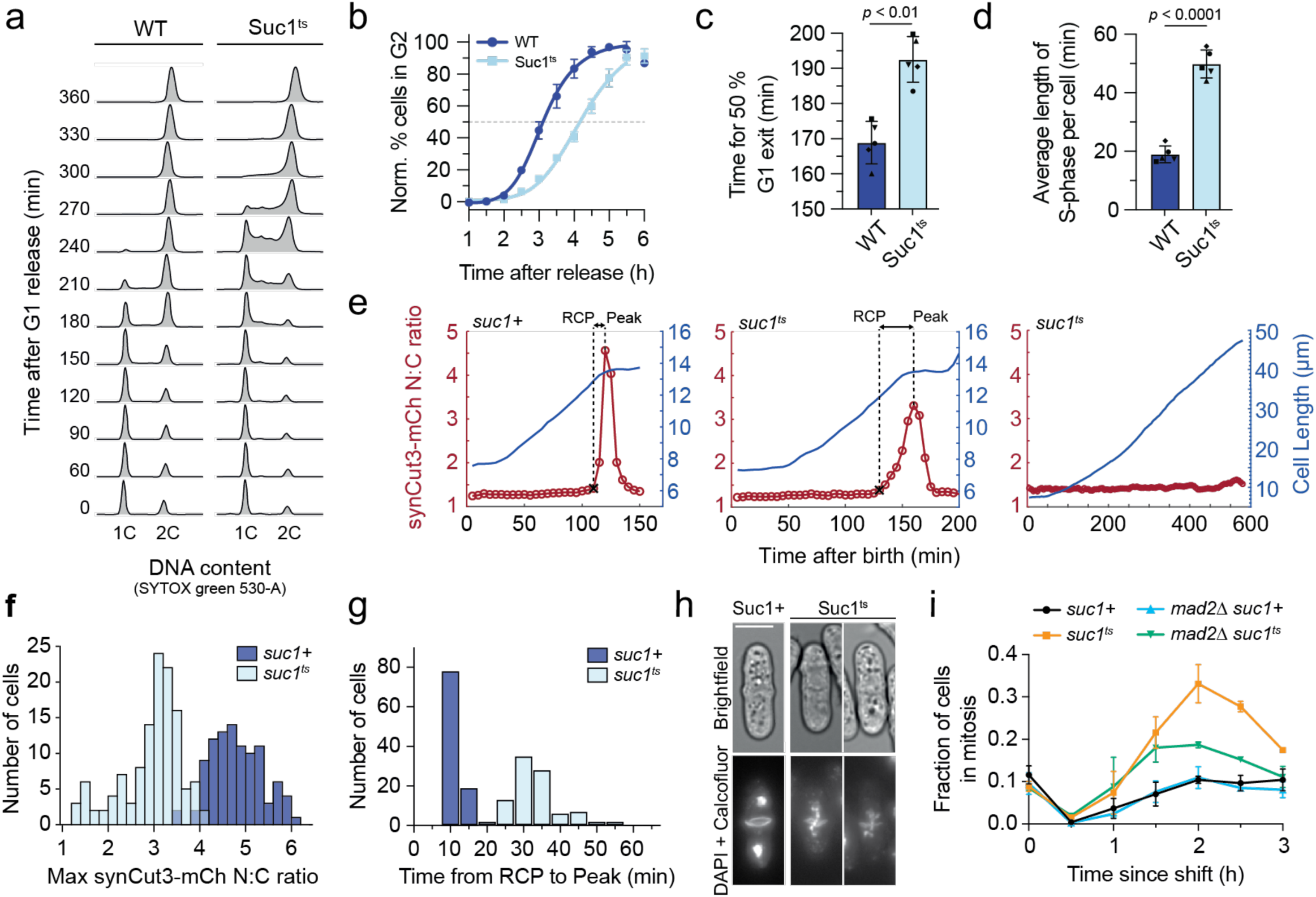
Suc1 regulates the onset and progression of both S-phase and mitosis. **a**, Representative histograms of DNA staining intensity via flow cytometry over time after release from a G1 arrest. **b**, Quantification of data in **a**, indicating the percentage of cells with 2C DNA content over time, normalised to time 0 min (Methods). Least-squares regression fit of a sigmoidal function. Mean and s.d. shown for *n* = 5 biological repeats. **c**, Time taken for 50 % of cells to exit G1, as determined by the 50 % point of sigmoidal curve fits (see also Extended Data Fig. 3b). Mean and s.d. shown for *n* = 5 biological repeats (denoted by shapes). **d**, Approximation of the mean time taken to complete bulk DNA replication per cell, as determined by the integration of the percentage of cells in S-phase over time (Methods). Mean and s.d. shown for *n* = 5 biological repeats (denoted by shapes). **e**, Representative single cell traces of cell length (blue) and CDK activity, as reported by synCut3-mCh sensor N:C ratio, (red) over time. RCP – rate change point. **f**, Maximum synCut3-mCh N:C ratio reached per cell trace. *n* = 98 cells (Suc1+), 124 cells (Suc1^ts^); representative of *n* = 2 biological repeats. **g**, Time between the CDK activity rate change point (RCP) and peak sensor N:C ratio. *n* = 98 cells (Suc1+), 95 cells (Suc1^ts^); representative of *n* = 2 biological repeats. **h**, Example images of heat-fixed cells 2 h after temperature shift, stained for DNA (DAPI) and division septa (Calcofluor). **i**, Fraction of cells with synCut3-mCh N:C ratio > 1.6 following temperature shift. *n* > 300 cells per timepoint, Mean and s.d. shown for *n* = 3 biological repeats (time 1.5 h, 2.5 h *n* = 2 biological repeats, time 0.5 h *n* = 1 biological repeat).

The G2/M transition was monitored using time-lapse microscopy of the CDK activity sensor synCut3-mCh^48^ (Extended Data Fig. 3c). Wildtype cells exhibit a switch-like increase in CDK activity as a result of mitotic CDK activation at the G2/M transition, triggering synCut3 nuclear import (Fig. 3e). In the *suc1^ts^* mutant ∼15 % of cells never underwent this CDK activation and continued growing without importing synCut3 or initiating mitosis (Fig. 3e, Extended Data Fig. 3d,e). The remaining cells displayed impaired CDK activation, with a 40 % reduction in peak sensor activity, and less switch-like CDK activation kinetics, taking three times longer than the wildtype to reach peak CDK activity (Fig. 3e-g). These cells also frequently became arrested in mitosis and exhibited defective nuclear segregation and subsequent bisection by the formation of one or multiple division septa (Fig. 3h, Extended Data Fig. 3a,d,e,f).

Suc1-dependent substrates at G2/M included proteins involved in centromere localisation, kinetochore attachment and spindle elongation (Fig. 2f, Extended Data Fig. 2i), so we investigated if the mitotic arrest phenotype of *suc1^ts^* was due to Spindle Assembly Checkpoint (SAC) activation^49^. SAC signalling is inactivated in *mad2Δ* cells^50^, which reduced the accumulation of mitotically arrested cells after Suc1 inactivation by 50 % (Fig. 3i). This indicates that loss of Suc1 function promotes SAC-mediated mitotic arrest in a significant fraction of cells. We conclude that Suc1 function is required both for mitotic onset and for multiple steps in mitosis, including involvement in the SAC.

To ensure the effects observed at G2/M were not confounded by defects earlier in the cell cycle, we synchronised cells in G2 with an ATP analogue-sensitive *cdc2^as^* allele before inactivation of Suc1. In the absence of Suc1 function, progression through mitosis was either delayed or blocked, and a substantial fraction cells displayed chromosome segregation and septation defects (Extended Data Fig. 3g). Overall these experiments demonstrate a direct role for Suc1 at G2/M and in mitosis.

We conclude that Suc1 has roles at the G1/S transition, in S-phase progression, in the activation of CDK at G2/M, and for mitotic progression. Therefore, the diversity of cell cycle phenotypes reported previously upon CKS inhibition has been recapitulated, but importantly, in a single model system. We propose that the changes in CDK substrate phosphorylation we have observed are responsible for these diverse cell cycle phenotypes.

### Suc1 drives phosphorylation of low affinity phosphosites by enhancing CDK activity

We next investigated the molecular basis of Suc1-dependent phosphorylation. No differences based on substrate sub-cellular localisation were detected (Extended Data Fig. 4a), indicating that Suc1 targets substrates in all sub-cellular compartments. We also found no difference in the pairwise comparison of phosphorylation of sites on the same protein or sites on different proteins, demonstrating that Suc1-dependence is not determined at the level of the protein substrate, but rather individual phosphosites (Extended Data Fig. 4b).

We compared the amino acid sequence surrounding each phosphosite and found enrichment for alanine residues in the +1 position of Suc1-dependent sites (Fig. 4a, Extended Data Fig. 4c). Although a +1 proline residue has been considered the minimal consensus sequence for CDK phosphorylation^51^, CDK has been reported to phosphorylate non-proline directed sites^41,52,53^, and several such non-canonical phosphosites have biological functions^35,36,54^. In our *suc1+* data, we observed phosphorylation of non-canonical CDK phosphosites at G2/M, and enrichment of these sites in the strong Suc1-dependent cluster (Fig. 4b). On average, non-canonical sites in *suc1^ts^* were phosphorylated to only 38 % of *suc1+* levels, compared to 65 % for proline-directed canonical sites (Fig. 4c). Non-canonical sites displayed a high frequency of basic residues in the +3 to +6 positions (Extended Data Fig. 4d), which increase CDK affinity^52^, consistent with previous *in vitro* reports^41,53^. No phosphorylation of non-canonical CDK sites was detected at G1/S. We conclude that Suc1 promotes CDK phosphorylation of non-consensus phosphosites lacking a +1 proline in mitosis *in vivo*.

**Fig. 4:**
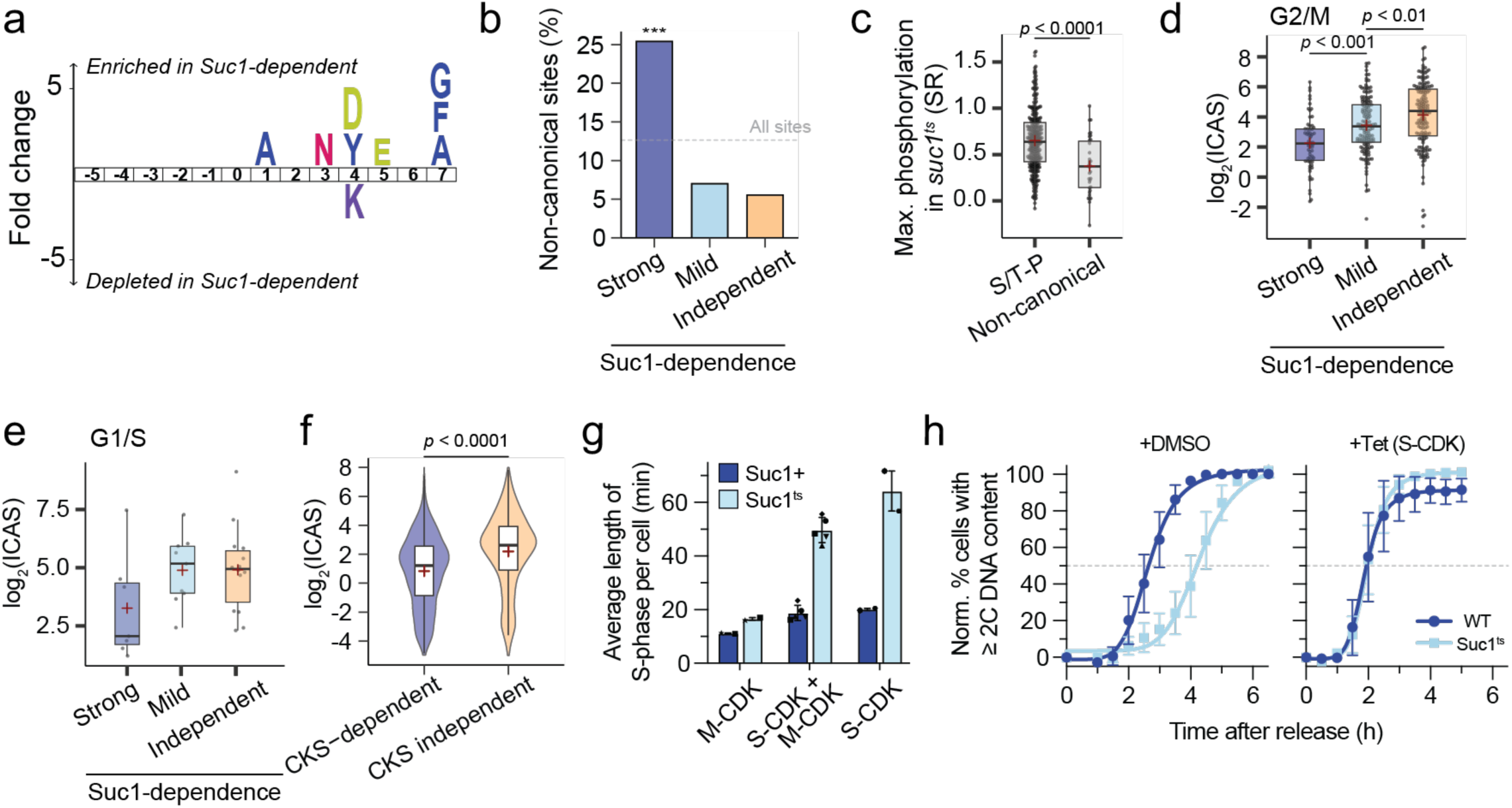
CKS enhances phosphorylation of intrinsically low-affinity phosphosites. **a**, IceLogo representation of amino acids enriched or depleted (*p* < 0.05) amongst Strong Suc1-dependent sites in the G2/M dataset, relative to Suc1 independent sites. **b**, Percentage of non-canonical phosphosites (lacking a +1 proline) in each behaviour class at G2/M. Dashed line indicates the frequency across the entire dataset. **c**, Maximum phosphorylation reached in the Suc1^ts^ condition, grouped by phosphosite type. *n* = 317, 34 for S/TP and non-canonical respectively. SR = Scaled ratio. 2 datapoints lie beyond the axis limits. **d** and **e**, ICAS for G2/M (**d**) and G1/S (**e**) phosphosites grouped by Suc1-dependence. Box plots represent IQR, with a black line at the median and a red + at the mean. *n* = 63, 143 and 145 for Strong, Mild and Suc1-independent respectively at G2/M; 1 data point is beyond the axis limits. *n* = 7, 9 and 14 for Strong, Mild and Suc1-independent respectively at G1/S. **f**, ICAS for *in vitro* phosphorylation data from ^41^ grouped by CKS-dependence, as determined in that study. Box plots represent IQR, with a black line at the median and a red + at the mean. *n* = 1110, 1653 for CKS-dependent and independent groups, respectively. 14 data points lie outside the axes bounds. **g**, Approximation of the mean time taken to complete bulk DNA replication per cell when S-phase is driven by different Cyclin-CDK complexes, as determined by the integration of the percentage of cells in S-phase over time (Methods). S-phase was driven by M-CDK via deletion of the other cyclins (*cig1Δ cig2Δ puc1Δ*), or S-CDK via transcriptional repression of *cdc13* (*P.nmt41-cdc13* thiamine repressible promoter). Note the data for S-CDK+M-CDK (WT) is the same as in Fig. 3d. Mean and range shown for *n* = 2 biological repeats of S-CDK and M-CDK conditions. Shapes denote corresponding repeats. **h**, Normalised percentage of cells with ≥2C DNA content over time after release from a G1 arrest (Methods), in the presence of only S-CDK activity (*P.nmt41:cdc13* + thiamine). Least-squares regression fit of a sigmoidal function. Tetracycline (Tet) addition results in over-expression of a Cig2-L-Cdc2 S-CDK fusion protein from a tetracycline-inducible promoter. ≤ 2C DNA content is shown instead of G2 population to account for S-CDK mediated endoreduplication cycles in the absence of M-CDK activity^91^. Mean and s.d. shown for *n* = 3 biological repeats.

We hypothesised that if Suc1 is required for phosphorylation of non-canonical sites due to their low affinity for CDK, this may generalise to proline-directed CDK sites which contain proximal unfavourable residues. Consistent with this, we observed depletion of favourable lysine residues and enrichment of unfavourable acidic residues downstream of Suc1-dependent phosphosites (Fig. 4a, Extended Data Fig. 4c). To quantify this, we employed a metric based on the *in vitro* substrate specificity of human CDK1^55^, which used the relative phosphorylation of peptide arrays by purified kinases to determine amino acid “preference scores” for each position around a phosphosite (Extended Data Fig. 4e). We applied these preference scores to our phosphosites yielding an “Intrinsic <u>C</u>DK <u>A</u>ffinity <u>S</u>core (ICAS)” (Methods). This reflects the efficiency of phosphorylation due to the peptide sequence in the absence of extrinsic factors such as docking interactions. We validated this metric using data from *Xenopus* Cyclin B1-CDK1 phosphorylation assays^52^, which revealed a strong correlation between *in vitro* phosphorylation of peptides and their ICAS (Extended Data Fig. 4f).

Suc1-dependent sites had lower ICAS values than Suc1 independent sites at both G2/M and G1/S (Fig. 4d,e). These differences persisted even when non-proline directed CDK sites were excluded (Extended Data Fig. 4g). We investigated if Suc1-dependent, low affinity sites might be phosphorylated only late in mitosis, when CDK activity is highest. However, there was no correlation between Suc1-dependency, or ICAS, and timing of phosphorylation during the cell cycle^4^ (Extended Data Fig. 4h,i). To test the generality of this effect, we also analysed phosphoproteomic data from *in vitro* Cyclin B-CDK1 kinase reactions performed with or without CKS1 in formaldehyde-fixed human TK6 cells^41^. CKS-dependent phosphosites in this study also had lower ICAS values on average than CKS independent phosphosites (Fig. 4f, Extended Data Fig. 4j). We conclude that CKS proteins promote the phosphorylation of intrinsically low affinity CDK phosphosites.

We hypothesised that dependence on Suc1 to phosphorylate weak CDK phosphosites might be overcome by increasing overall CDK activity. The analysis of S-phase progression in the absence of Suc1 was therefore repeated using genetic backgrounds in which CDK activity results solely from S-CDK (Cig2-Cdc2) activity, or M-CDK (Cdc13-Cdc2) activity. S-phase driven by M-CDK suppressed the slow S-phase phenotype in the absence of Suc1 function, whereas S-phase driven by S-CDK was slower than in a wildtype background (Fig. 4g, Extended Data Fig. 4k). The slow S-phase phenotype in the absence of Suc1 function was also rescued by overexpression of an S-CDK fusion protein in the *suc1^ts^* strain (Fig. 4h, Extended Data Fig. 4l), demonstrating that it is increased CDK activity, and not the identity of the cyclin which compensates for loss of Suc1 function at G1/S. We conclude that Suc1 enhances CDK activity to drive phosphorylation of a subset of intrinsically weak CDK phosphosites.

### Suc1 enhances local CDK activity to promote multisite phosphorylation *in vivo*

The binding of CKS proteins to ‘priming’ phosphosites via the conserved phosphate-binding pocket has been reported to drive multisite phosphorylation of a substrate^36,37^ (Extended Data Fig. 5a). Such binding could act as a mechanism to enhance CDK activity by stabilising substrate-CDK complexes, thus promoting phosphorylation of nearby low-affinity sites. We compared phosphosites on singly and multiply phosphorylated substrates, and found that phosphosites on multiply phosphorylated proteins had lower ICAS values and were phosphorylated less well in the absence of Suc1 function (Fig. 5a, Extended Data Fig. 5b,c). In addition, two phosphate-binding pocket mutant alleles of *suc1* could not rescue the lethality of *suc1^ts^*, demonstrating that phosphate binding is essential for at least some Suc1 function(s) (Extended Data Fig. 5d).

**Fig. 5:**
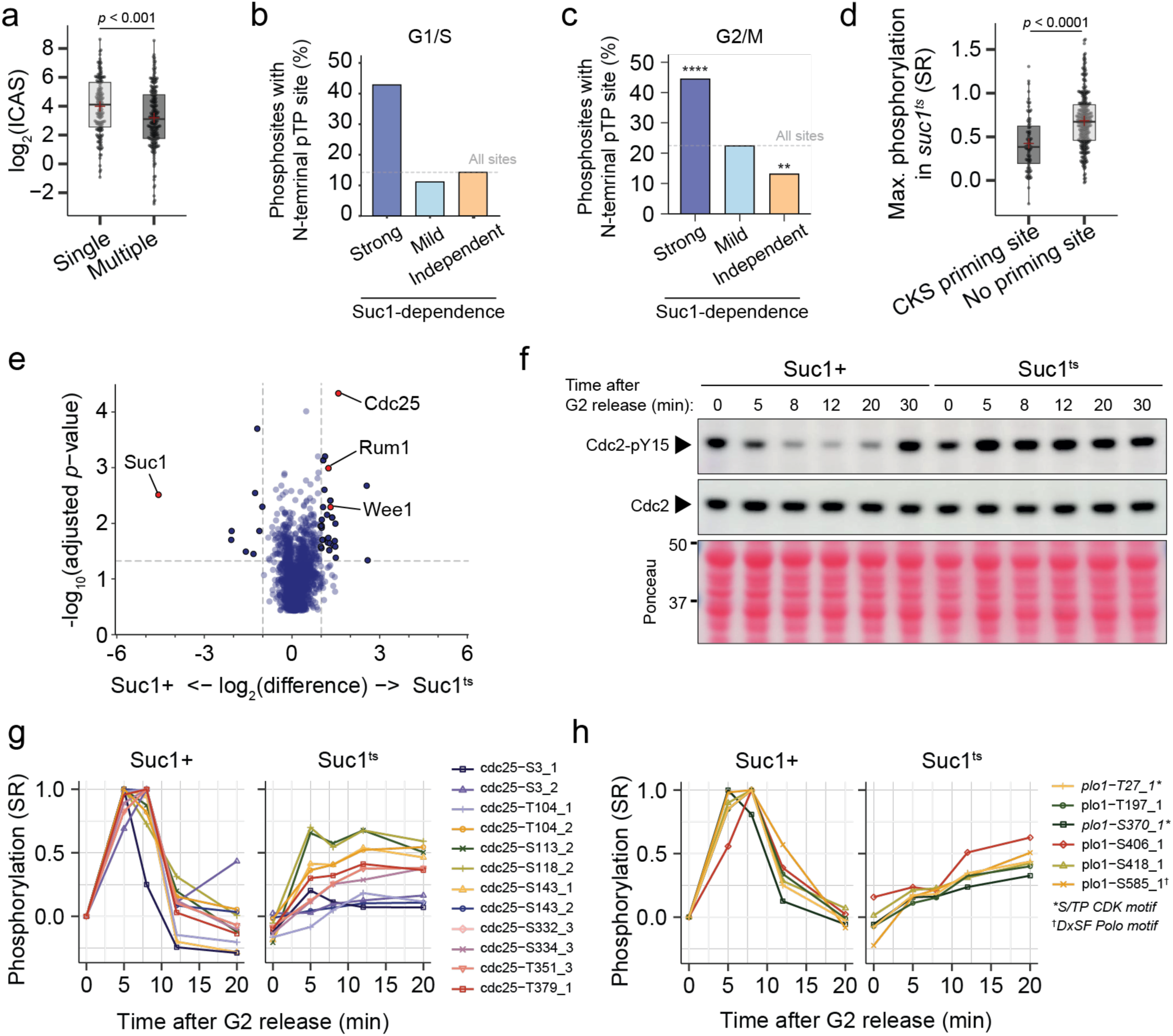
Suc1 enhances local and global CDK activity. **a**, ICAS for G2/M phosphosites grouped by how many CDK sites have been identified on the protein substrate (Single vs Multiple). Box plots represent IQR, with a black line at the median and a red + at the mean. *n* = 151, 200 for single and multiple respectively. 2 data points are beyond the axis limits. **b** and **c**, Percentage of G1/S (**b**) and G2/M (**c**) phosphosites in each behaviour class with a putative upstream pT-P CKS-priming phosphosite. Dashed line indicated the frequency across the entire dataset. **d**, Maximum phosphorylation reached in the Suc1^ts^ condition at G2/M, grouped by the presence or absence of a putative upstream pT-P CKS priming phosphosite. Box plots represent IQR, with a black line at the median and a red + at the mean. SR = Scaled ratio. 8 datapoints lie beyond the axis limits. *n* = 79, 272 for CKS priming site and no site, respectively. **e**, Multiple testing corrected T-test significance (Q values) vs fold change in protein abundance detected by mass spectrometry following Cdc2-GFP immunoprecipitation. *n* = 3 biological replicates per strain. Note this dataset is the same as Fig. 1f. **f**, Western blot for Cdc2 protein and Cdc2-Y15 phosphorylation after release from a G2 arrest at the restrictive temperature. Representative of 3 biological repeats. For gel source data, see Supplementary Fig. 1. **g** and **h**, Selected CDK phosphosites on Cdc25 (**g**) and Plo1 (**h**) during the G2/M time course. _1,_2,_3 represents the multiplicity state of the phosphopeptide detected. SR = scaled ratio.

To test if Suc1 docking to ‘priming’ residues could explain Suc1-dependent phosphorylation, we analysed the prevalence of putative CKS-priming sites containing a phospho-threonine 12-52 amino acids N-terminal of each phosphosite, in accordance with reported criteria^37,38^. Phosphosites with putative pT-P priming sites upstream were enriched among strong Suc1-dependent phosphosites at G1/S and G2/M, and displayed lower average phosphorylation levels across all clusters compared to phosphosites without priming phosphorylation (Fig. 5b,c,d, Extended Data Fig. 5e). When grouping by distance to the putative priming site, we found stronger enrichment at distances of 13-18 amino acids upstream, suggesting the most effective priming occurs at these shorter distances (Extended Data Fig. 5f). No enrichment was observed for substrates with C-terminal pT-P phosphorylation (Extended Data Fig. 5g) nor priming on serine residues (Extended Data Fig. 5h,i,j). This supports the N-C directionality and phospho-threonine specificity of CKS-mediated priming^37,38^, although only 40 % of Strong Suc1-dependent sites have putative priming sites (Fig. 5b,c). This indicates that priming on N-terminal pT-P sites does not explain the majority of Suc1-dependent phosphorylation *in vivo*.

### Suc1 also regulates global CDK activity

We broadened our analysis to investigate if Suc1 regulates global CDK activity in addition to its substrate-specific effects, by examining the interaction of CDK regulators in the presence or absence of Suc1 using Cdc2 co-immunoprecipitation data. This revealed enhanced association of the CDK inhibitor Rum1 with Cdc2 in the absence of Suc1 function (Fig. 5e). Rum1 inhibits Cdc2 to regulate the G1/S transition^56^, suggesting that enhanced Rum1 association in the absence of Suc1 function may decrease global CDK activity and contribute to the defects in G1/S progression. But most notably, we observed a 2.5-fold increase in the association of the CDK inhibiting kinase Wee1, and a 3-fold increased association of the CDK-activating phosphatase Cdc25 with Cdc2 in the absence of Suc1 function (Fig. 5e). Since CDK activity at the G2/M transition is primarily regulated by phosphorylation of Cdc2-Y15 by Wee1 and dephosphorylation by Cdc25^57–60^, we tested the impact of Suc1^ts^ on Cdc2-Y15 phosphorylation. In *suc1+* cells, Cdc2-Y15 phosphorylation levels rapidly decreased during the transition from G2 into mitosis, but in the absence of Suc1 function Cdc2-pY15 was not dephosphorylated (Fig. 5f, Extended Data Fig. 5k). This is consistent with *suc1^ts^* cells failing to properly activate CDK (Fig. 3e,g) and confirms that Suc1 regulates Cdc2-Y15 phosphorylation. Enhanced binding of Cdc25 to CDK should promote Cdc2-pY15 dephosphorylation, suggesting that the enhanced association of Wee1 dominates that of Cdc25. No differences were observed in Cdc2-Y15 phosphorylation at the G1/S transition (Extended Data Fig. 5l).

CDK phosphorylates Cdc25 and Wee1 to regulate their enzymatic activities, resulting in feedback loops that amplify CDK activation at the G2/M transition^61,62^. The CDK-dependent activating phosphosites on *S. pombe* Cdc25 have been mapped^63^, so we also examined the effect that Suc1 has on CDK phosphorylation of these sites. In the absence of Suc1 activity, we observed substantial hypophosphorylation of 9 activating phosphosites on Cdc25 (Fig. 5g). Therefore, Suc1 also modulates the feedback phosphorylation of Cdc25, consistent with the impaired Cdc2-pY15 dephosphorylation we observed. Polo kinase also contributes to the CDK activation, and although specific phosphosites have yet to be mapped, mitotic phosphorylation of Polo enhances its activity^64^. Loss of Suc1 also impairs phosphorylation of multiple phosphosites on Plo1 (Fig. 5h). We conclude that Suc1 plays a critical role in the feedback control regulating global CDK activity via CDK-Y15 phosphorylation at the G2/M transition, through modulating CDK interactions with Cdc25 and Wee1, the phosphorylation of Cdc25, and probably also by impacting Polo.

## Discussion

We have shown that loss of CKS (Suc1) function in fission yeast cells delays the G1/S transition, slows down S-phase progression, impairs the G2/M transition and blocks mitotic progression. This recapitulates in one eukaryotic model system the diverse phenotypes reported independently from multiple organisms and cell types^9,13,15–24^, establishing that they are not the consequence of divergent roles for CKS proteins in different organisms. It also highlights the critical role of CKS proteins throughout the cell cycle.

We found that loss of CKS function results in widespread impaired phosphorylation of 221 of 390 CDK phosphosites on 133 protein substrates *in vivo*, 51 of which have been implicated in cell cycle functions. These include the G2/M regulators Polo kinase, INCENP, CDC14, and the CDK-activating phosphatase Cdc25, as well as proteins involved in kinetochore attachment, chromosome condensation and segregation, and the SAC. CKS-dependent substrates also act at G1/S, including 5 CKS-dependent sites on the essential origin firing factor Drc1^Sld2^ ^47^. These results establish that CKS modulates CDK substrate phosphorylation throughout the entire cell cycle. Given that human CKS1B or CKS2 can substitute for Suc1, and widespread CKS-dependent phosphorylation of the human proteome *in vitro,* including DNA replication proteins^41^, it can be expected that CKS-dependent phosphorylation will be extensive in mammalian cells *in vivo*. This broadens the role of CKS at the G1/S transition, which has previously been thought to act only via the degradation of CDK inhibitors p21 and p27 in metazoa^16,30,31^. Supporting this, a p27-independent effect of CKS1 at G1/S in mammalian cells has been observed^18^. We conclude that the majority of the diverse impacts of CKS proteins on the cell cycle can be understood based on widespread changes in the phosphorylation of large numbers of CDK substrates involved in diverse cell cycle processes.

We found that CKS is required to enhance the phosphorylation of intrinsically weak CDK phosphosites *in vivo*, including non-proline directed sites. This is supported by human *in vitro* phosphorylation data reporting CKS-enhanced CDK phosphorylation of non-proline directed sites^41^, assays of individual reporter constructs in budding yeast^38,65^, and by our finding here that increasing CDK activity at G1/S *in vivo* rescues the S-phase defects in the absence of CKS function. Our data also indicate that for a given phosphosite, different Cyclin-CDK complexes differ in their requirement for CKS, likely due to differences in CDK activity, since M-CDK is not dependent on Suc1 to drive a normal S-phase, but S-CDK is. Therefore, we propose that a conserved, general role of CKS proteins is to enhance CDK phosphorylation of intrinsically weak phosphosites.

Why should regulatory proteins such as CKS be required to drive phosphorylation of poor CDK substrates? One proposal has been that low affinity substrates may broaden the range of phosphorylation timings during the cell cycle, only becoming phosphorylated at very high CDK activity^54^. However, we found no correlation between intrinsic phosphosite sensitivity and the timing of phosphorylation during the cell cycle. As an alternative, we propose that the requirement for CKS proteins is the result of CDK having hundreds of protein substrates which have disparate biochemical functions. Individual phosphosites may be evolutionarily constrained to maintain both affinity for the CDK active site and for binding other molecules necessary to perform their functions. CKS proteins could relax these constraints, allowing phosphorylation of sub-optimal sites, thus extending the number of proteins subject to CDK control. Docking to the hydrophobic patch or the phosphate-binding pocket of cyclins, perhaps in conjunction with CKS interactions, may also alleviate this constraint^39,66^. This principle would be particularly important for cell cycle control, given the number of events that require co-ordination and the complexity of their function.

How does CKS enhance the activity of the Cyclin-CDK complex to drive phosphorylation of weak phosphosites? Our *in vivo* studies demonstrated that Suc1 regulates the association of Cdc25 and Wee1 with CDK and the phosphorylation of Cdc25 and Polo kinase, promoting Cdc2-Y15 dephosphorylation and thus increasing global CDK activity. *In vitro* structural studies suggest that the binding of human CKS1 and CDC25A to CDK2 are mutually exclusive based on overlapping CDK interaction interfaces^67^, indicating CKS may regulate CDC25C and CDK1 association in an analogous competitive manner. CKS promotes Cdc2-Y15 dephosphorylation in *Xenopus* extracts^9,68^, and Wee1 and Cdc25 phosphorylation *in vitro*^68,69^. Therefore, the regulation of CDK-Y15 phosphorylation by CKS appears to be conserved across eukaryotes.

We found Suc1 also regulates the association of CDK with the G1/S inhibitor Rum1, which may further regulate global CDK activity. CKS proteins have been shown to regulate G1/S CDK inhibitors in other organisms by promoting their degradation^16,30,31,36^, however our data suggest CKS also regulates the association of CDK with CDK inhibitor proteins *in vivo*, consistent with *in vitro* observations^70^.

Our demonstration that CKS inhibits the binding of Cdc25 and Wee1 to the Cyclin-CDK complex *in vivo* has implications for the Cdc2-Y15 phosphorylation/dephosphorylation futile cycle. It indicates that the association of a CKS protein with the Cyclin-CDK complex prevents access of Cdc25 and Wee1 to Y15, temporarily stabilising the phosphorylation status and activity of CDK, until CKS protein dissociation. The association and dissociation of CKS therefore would provide a window for substrate phosphorylation, in which CDK is not subject to Cdc2-pY15 turnover by Cdc25 and Wee1, separating CDK catalysis and Y15 phospho-regulation in a CKS-dependent manner.

However, downregulation of CDK activity by Y15 phosphorylation does not explain all the effects of Suc1 inactivation on CDK substrate phosphorylation. Firstly, CKS-dependent CDK substrate phosphorylation also occurred at G1/S, where loss of Suc1 function does not impact Cdc2-Y15 phosphorylation. Secondly, the correlation between CKS dependence and intrinsic substrate sensitivity persists in our analysis of *in vitro* data which is not subject to regulation by Y15 phosphorylation. Thirdly, a CDK-Y15F mutant bypassing Y15 regulation rescues mitotic entry but not mitotic arrest after CKS depletion in *Xenopus* extracts^9^. It has been proposed that the presence of CKS bound to CDK also enhances basal kinase activity, based on *in vitro* kinase assays against a *S. cerevisiae* Cdc16 peptide^66^. We propose a variation on this, that CKS proteins enhance Cyclin-CDK activity towards specific substrates by stabilising their association with the CDK complex. This could involve binding to nearby phosphorylated residues given the essentiality of the CKS phosphate-binding pocket, and our finding of the enrichment of CKS-dependent and intrinsically weak phosphosites on multiply phosphorylated proteins. An N- to C-terminal CKS-mediated multisite phosphorylation mechanism likely contributes to this^36,37^, although we have shown this can only account for around 40 % of Suc1-dependent sites *in vivo*. Multiple stabilisation modes may therefore exist, including some that are independent of priming phosphorylation and the CKS phosphate-binding pocket^71^.

We conclude that CKS controls CDK protein kinase activity globally by modulating its association with Wee1 and Cdc25 thus regulating CDK-Y15 phosphorylation, and also propose modulation occurs locally through stabilising CDK-substrate interactions. CKS thereby enhances the phosphorylation of around 50 % of CDK phosphosites, particularly intrinsically weak sites, impacting the onset and progression through S-phase and mitosis. This establishes CKS as major regulatory component of Cyclin-CDK complexes, controlling progression through the entire cell cycle.

## Methods

### Schizosaccharomyces pombe genetics and cell culture

Fission yeast strains were constructed by genetic crosses or lithium acetate transformation as described previously^72^. Strain genotypes were checked by colony PCR, and DNA sequencing where necessary. All strains used for this study and the plasmids required for their constructions are listed in (Supplementary Table 4). Growth media and conditions were as described previously^72^, with minor modifications. Strains were incubated at 22 °C rather than 25 °C when grown on solid agar media and inoculated into liquid media as small colonies rather than dense patches, to minimise diploidisation. Strains with the *cdc2(as)* allele were cultured in media supplemented with 1 M D-sorbitol. Unless specified, all experiments were carried out at 25 °C as the permissive temperature in yeast extract medium, supplemented with adenine, leucine, histidine, and uracil to 0.15 g/L, and shifted to 36 °C as the restrictive temperature. Experiments involving thiamine-repressible promoters were conducted in Edinburgh Minimal Medium (EMM2, MP Biomedicals), supplemented with leucine to 0.15 g/L for auxotrophic strains. Cells were maintained in exponential growth (2-10 x 10^6^ cells/mL) for at least 36 hours before all experiments. Thiamine hydrochloride (Sigma) in water was added to a concentration of 30 µM to repress thiamine-repressible promoters. Anhydrotetracycline (Sigma) dissolved in DMSO was added to a concentration of 0.3125 µg/ml to induce expression from tetracycline-inducible promoters. 1-NMPP1 (Toronto Research Chemicals) dissolved in DMSO was added at a concentration of 1 µM to inhibit the *cdc2(as)* allele for G2 arrests. 1-NMPP1 blocks were released by washing three times with pre-warmed media via filtration. The total DMSO volume never exceeded 0.1% v/v to avoid DMSO-mediated toxicity.

To arrest cells by nitrogen starvation, cells were collected by centrifugation (1,800 *g*, 2 min), washed four times with 50 mL EMM2 media lacking any nitrogen source, resuspended at a density of 2 x 10^6^ cells/mL and incubated for 16.5 h at 25 °C with shaking. Where strains were auxotrophic for leucine, the arrest media was supplemented with 0.05 g/mL leucine. After 16.5 h cells were shifted to 36 °C for 1 h, collected by centrifugation (1,800 *g*, 2 min) or filtration, and resuspended in pre-warmed EMM2 media. Where thiamine-repression was required, thiamine hydrochloride was added at the same time as the temperature shift. Where tetracycline-induced expression was required, tetracycline was added immediately after release into nitrogen containing media.

### Generation of the *suc1^ts^* allele

The *suc1-2* temperature sensitive allele (*suc1^ts^*) was generated by targeted *in vitro* mutagenesis. A synthetic DNA fragment encoding the analogous mutations to the budding yeast *CKS1-ts38* allele in *suc1*, suc1(A32T,E35Y,N57D,L74H):BsdMx6 (Integrated DNA Technologies), was amplified by error-prone PCR using Taq DNA polymerase (New England BioLabs) for 20 cycles. 200 reactions were performed individually then pooled. This DNA was then subjected to mutagenesis by incubation with 1 M hydroxylamine hydrochloride pH 6.7 (with 5 M NaOH) for 1 h at 70 °C. The DNA was purified using a Qiagen spin column, split into 100 fractions, and transformed into wild type cells. Cells were recovered at 25 °C before replica plating to 36 °C to screen for temperature-sensitivity.

### Microscopy

Microscopy was performed using a Nikon Ti2 inverted widefield microscope with an Okolab environmental chamber, a 100X Plan Apochromat oil-immersion objective (NA 1.45), a perfect focus system (PFS) and a Prime sCMOS camera (Photometrics). Fluorescence imaging was performed using a SpectraX LED light source (Lumencor). mNeonGreen was imaged by excitation at 470/24 nm (with a Chroma ET525/50m single-band bandpass emission filter); mCherry at 575/25 nm (with a Semrock FF02-641/75-25 single-band bandpass emission filter) and DAPI/Calcofluor at 395/25 nm (with a Semrock FF02-438/24-25 single-band bandpass emission filter). Five z-slices were acquired; −1 µm to +1 µm, at 0.5 µm interval. This was controlled with MicroManager (µManager, v2.0) software^73^ (Open-imaging). Timelapse microscopy samples were imaged on 1 % agarose media pads^74^ with the chamber set to 36 °C.

Raw images were processed using ImageJ^75^, and custom MATLAB scripts. Cell segmentation was carried out from brightfield images using YeaZ^76^ or custom MATLAB scripts^77^, and binary cell masks manually corrected. Cell length was determined using the maximum Feret diameter of cell masks. Fluorescence images were maximum z-projected, and whole cell intensities measured as the mean pixel value of each cell mask. mNeonGreen intensities were background subtracted by the mean green fluorescence intensity of an untagged control strain. Nuclear to cytoplasmic ratio was estimated by ratio of the mean of the top 15% of pixels to the bottom 85% of pixels, since the nucleus occupies approximately 15% of the cross-sectional area of a *S. pombe* cell. Single cell timelapse traces were generated as described^77^.

### S-phase progression and DNA content analysis

To measure DNA content, 520 µL of culture were fixed by addition to 1.4 mL of ice-cold 96% ethanol, resulting in a 70% v/v ethanol suspension. After >24 h at 4 °C, cells were collected (800 *g*, 3 min), washed once with 700 µL 50 mM sodium citrate before resuspension in 500 µL 50 mM sodium citrate containing 0.1 mg/mL RNaseA (Sigma) for 3 - 18 h (37 °C, 1000 rpm). 200 µL 50 mM sodium citrate containing 3.5 µM SYTOX Green (Thermo Fisher) was added, and samples were vortexed for 1 min before sonication for 10 s using a probe sonicator (Soniprep 150 plus) at amplitude 3. >10,000 cells were analysed per sample on a BD LSRFortessa cell analyser. DNA content (530/30-A) is presented on a linear scale after gating for single cells in FlowJo (v10.9, Treestar Inc.) based on FSC-H v FSC-A then SSC-A v FSC-A profiles.

The percentage of cells in G1, S and G2 after release from nitrogen-arrest were determined by gating 595-A histograms. 1C and 2C populations were gated symmetrically around their peaks, with cells between these two gates deemed to be in S phase. To account for differences in arrest efficiency, the percentage of cells in G1 (%G1) at each timepoint was normalised by dividing by the %G1 at time 0 (except one repeat of the *cig1Δcig2Δpuc1Δ* experiment in which time 0 was lost and time 60 was used). The %S phase or %G2 were normalised by subtracting the %S phase or %G2 at time 0 and then dividing by the %G1 at time 0. 50 % G1/S transition times were determined by the IC_50_ of a least-squares fitting of a sigmoidal curve to the normalised %G1. The average length of S phase was approximated by integrating the normalised %S phase over time using the composite trapezoidal rule, then dividing by the percentage of cells which had undergone G1/S at the final time point (100 - %G1). If the normalised %S decreased below −1% only values up to that time point were included in the calculation. *cig1Δcig2Δpuc1Δ* were integrated up to 300 min to minimise the effects of the pronounced G1 population present in asynchronous populations of these cells on S-phase duration estimates. Due to the occasional failure of DNA staining some samples were lost and the *n* number of a specific timepoint is lower than the experiment. These are: Fig. 3a,b, Extended Data Fig. 3b – t=1h both strains n=4, t=6h Suc1+ n=4; Extended Data Fig. 4k – t=1.5h Suc1+ n=1; Extended Data Fig. 4k – t=0h both strains n=1; Fig. 4h, Extended Data Fig. 4l – t=1h all strains n=2, t=6.5h Suc1+ DMSO n=2.

### Determination of mitotic progression

To determine binucleation and septation indices, 2.5 µL of culture was heat fixed at 70 °C before addition of 4’,6-diamidino-2-phenylindole (DAPI, Invitrogen) to stain DNA and Calcofluor white (Sigma) to stain septa. Cells were considered binucleated if there was clear separation of the two nuclear masses, and septated once a complete septum could be detected. Cells with asymmetric DNA segregation, fragmented nuclei, mis-placed or multiple septa and septa bisecting the nuclear masses were considered aberrant.

### Serial dilution assays

Serial dilution assays were performed using cells in exponential growth. 200 µL of cultures at a density of 5 x 10^6^ cell/mL were 10-fold serially diluted in a 96-well plate. A sterilised 6 x 8 grid of metal pins was used to transfer approximately 4 µL to agar plates.

### Protein extraction

Protein extraction for western blots and G2/M phosphoproteomics was performed via trichloroacetic acid (TCA) precipitation. Cell cultures were quenched by addition to 100% ice-cold TCA to yield a final 10% v/v TCA concentration. After >30 min on ice cells were collected (3 min, 1,800 *g*), washed with 50 mL ice-cold 10% TCA, then with 1 mL ice-cold acetone, and dry pellets frozen at −80 °C. Cells were washed twice in 500 µL Beating Buffer (BB, 8 M urea, 50 mM ammonium bicarbonate, 5 mM EDTA, 1 mM phenylmethylsufonyl fluoride (PMSF), 1X cOmplete EDTA-free protease inhibitor cocktail (Roche), 1X phosSTOP phosphatase inhibitor cocktail (Roche)) before resuspension in 50 µL BB with 1 mL of 0.4 mm acid-washed glass beads (Sigma). Cells were ruptured by three cycles of mechanical beating at 4 °C in a FastPrep120 (2 x 30 s 5.5 m/s, 1 x 45 s 6.5 m/s). Cell debris was then pelleted by centrifugation at 4 °C (15 min, 16,000 *g*). The concentration of the supernatant was quantified using a Quick Start Bradford Protein Assay (BioRad) with BSA as a standard. Phosphoproteomics samples were additionally diluted by 1:1 v/v addition of 7 mM magnesium chloride, 0.02 U/µL Benzonase (Merck) in water and incubated at 37 °C for 1.5 h.

Protein extraction for G1/S phosphoproteomics was performed via a modified TCA precipitation protocol to improve recovery from nitrogen arrested cells. Cell cultures were rapidly quenched by addition to 100% ice-cold TCA to yield a final 10% v/v TCA concentration. After >30 min on ice, cells were collected (3 min, 1,800 *g*), washed in 1 ml ice-cold 20 % TCA, then 500 µl 12.5 % TCA, before resuspension in 100 µl 12.5 % TCA with 1 mL of 0.4 mm acid-washed glass beads (Sigma). Cells were ruptured by three cycles of mechanical beating at 4 °C in a FastPrep120 (3 x 45 s 5.5 m/s). Proteins were pelleted by centrifugation at 4 °C (20 min, 16,000 *g*), and resuspended thoroughly in 1 ml ice-cold acetone. Samples were centrifuged at 4 °C (5 min, 16,000 *g*), and the pellets dried for < 60 s in a Speedvac, before resuspension in 100 mM HEPES pH 7, 5 % SDS, 100 mM DTT, 1 mM EDTA, 1 mM PMSF, 1X cOmplete EDTA-free protease inhibitor cocktail (Roche), 1X phosSTOP phosphatase inhibitor cocktail (Roche). Samples were heated to 70 °C for 15 min, sonicated in a Bioruptor at maximum power for 15 min (30 s on/30 s off), and clarified by centrifugation (5 min, 16,000 *g*). The supernatant was quantified using a Pierce A660 Protein Assay (ThermoScientific) supplemented with Ionic Detergent Compatibility Reagent and with BSA as a standard.

Protein extraction for immunoprecipitation was performed via cryo-milling. 750 mL of culture was pelleted by centrifugation (3 min, 2,000 *g*), and the pellet resuspended in 1 mL lysis buffer (50 mM HEPES pH 7.5, 75 mM KCl, 1 mM MgCl_2_, 0.1 % NP-40, 1X cOmplete EDTA-free protease inhibitor cocktail (Roche), 1X phosSTOP phosphatase inhibitor cocktail (Roche)). Cells were frozen in small pellets by drop-wise addition into liquid nitrogen. Pellets were ground in a CertiPrep 6850 Freezer/Mill for 3x 2 min cycles (15 CPS), and the powder stored at −80 °C. Aliquots were resuspended in 50 µL ice-cold IP buffer (50 mM Tris-HCl pH 7.5, 100 mM KCl, 2 mM MgCl_2_, 0.2 % Triton-X, 1X cOmplete EDTA-free protease inhibitor cocktail (Roche), 1X phosSTOP phosphatase inhibitor cocktail (Roche)), sonicated using a Bioruptor (maximum power, 20 s), clarified by centrifugation (5 min, 16,000 *g*, 4 °C) and the supernatant recovered.

### Western blotting

30 µg of protein lysates were separated using denaturing polyacrylamide gel electrophoresis and transferred to 0.45 µm PVDF membranes via wet transfer at 30 V for 16 h. Total protein staining on membranes was performed with Ponceau S solution (Sigma). The following primary antibodies were used: anti-Cdc2 at 1:20,000 (Y63.2 mouse monoclonal, lab stocks), anti-Cdc2-pY15 at 1:1000 (#9111 rabbit polyclonal, Cell Signalling Technology). The following secondary antibodies were used: 1:5,000 horseradish peroxidase-conjugated donkey anti rabbit (NA934, GE Healthcare) or 1:5,000 goat anti-mouse (STAR120P, AbD SeroTEC). Signal was detected using SuperSignal West Femto Maximum Sensitivity Substrate (34095, Life Technologies) and imaged on an Amersham Imager 600.

### Co-immunoprecipitation mass spectrometry

Samples were incubated with 30 µL Protein G dynaBeads (ThermoFisher), which had been cross-linked to anti-GFP antibodies (11814460001, Roche) with dimethyl pimelimidate, for 45 min at 4 °C. Beads were washed 3x with 1 mL ice-cold IP buffer before elution in 30 µL 50 mM Tris 7.5, 5% SDS (1 min, 55 °C, 600 rpm). Samples were digested using S-trap (Protifi) and data were acquired using Evosep system coupled to a TIMS TOF Ultra using DIA PASEF. The Evosep system was filtered with an Ion Optics Aurora Elite column. Whisper Zoom 40SPD method was used for chromatography. Data were processed using Spectronaut 19 using the directDIA procedure with standard settings and an *S. pombe* Uniprot database.

### Tandem mass tag proteomics

200 µg of protein lysates per time point were reduced with 5 mM DTT (25 min, 56 °C) and alkylated with 10 mM iodoacetamide (30 min, room temperature, in the dark) before quenching with 7.5 M DTT. Samples were either digested using on-bead SP3 methodology^78^ (for G2/M experiment), or using S-Trap (Protifi, G1/S experiment), both with the variation that 50 mM HEPES (pH 8.5) was used in place of ammonium bicarbonate. Peptides were labelled with a TMT11plex (G2/M) or TMTpro16plex (G1/S) Isobaric Label Reagent Set (ThermoFisher) according to the manufacturer’s instructions. Samples were quenched with hydroxylamine and then mixed, before drying by vacuum centrifugation. Peptides were acidified and desalted using Pierce Peptide Desalting Columns (ThermoFisher), and initial phosphopeptide enrichment performed using a TiO2 phosphopeptide enrichment kit (ThermoFisher). Eluted peptides were dried and stored at −80 °C and a second round of enrichment with a Fe-NTA phosphopeptide enrichment kit (ThermoFisher) performed on the flow through. The pooled phosphopeptide enrichment fractions and 10% of the flow through fraction (proteome) were desalted and fractionated using a high pH reversed phase fractionation kit (ThermoFisher). The buffer series for native peptides was used for the phosphopeptide enriched samples (due to the physical properties conferred by the phosphate groups) and the buffer series for TMT-labelled proteins was used for the proteome fractions. Samples were analysed using an Orbitrap Eclipse Tribid mass spectrometer (ThermoFisher) coupled to an UltiMate 3000 HPLC system. Phospho-enriched fraction was injected in duplicate using a 3 h gradient from a 75 μm × 50 cm C18 column, (MS2 HD and MSAOTMS3 methods), proteome fractions were injected using 3 h gradients and analysed using the MS2 HCD (G2/M) or MSAOTMS3 (G1/S) methods.

### Mass spectrometry data analysis

Raw mass spectrometry data files were processed using MaxQuant (v1.6.14.0 (G2/M) and v2.4.10.0 (G1/S))^79^. A UniProt *S. pombe* reference proteome was used for searches, with an additional entry for the Suc1^ts^ protein. The default settings were used, with Phospho(STY) added as a variable modification for phospho-enriched samples. MaxQuant Phospho STY.txt (phosphopeptides) and ProteinGroup.txt (peptides) output files were loaded into Perseus (version 1.6.14.0)^80^, annotated and filtered to remove common contaminants and reversed entries. Proteome samples were also filtered to remove sites only identified by modified peptides. These matrices were then exported for further processing using custom Python and R scripts. Sites were filtered for valid values (100 %), localisation probability (> 0.75), and median normalised. Only multiplicity 1 peptides were analysed, except for Fig. 5g and h. Scaled ratios were calculated according to the formula (X – t0_Suc1+_)/(max_Suc1+_ – t0_Suc1+_) for G2/M, where X is the intensity of a given timepoint. For G1/S the variation (X – t90)/(max_Suc1+_ – t90) was implemented, using the respective time 90 min intensity as a baseline for each strain, to account for differences in arrest efficiency.

### Identification of CDK sites of interest

CDK sites of interest at G1/S and G2/M were determined by first filtering for phosphosites we have identified previously^4^ and in unpublished data, which are dephosphorylated by >50% within 12 min upon acute CDK inhibition. These sites were scaled as above and clustered by Euclidean distance. Clusters displaying the expected behaviour – increasing in phosphorylation at the relevant transitions were selected and further filtered to include only those with a 2-fold minimum fold change. These clusters included known G1/S (“Early”) and G2/M (“Late”) substrates^4^, validating their selection. See also Extended Data Fig. 2.

### CDK substrate analyses

To calculate ICAS values, the scaled values from the position-specific scoring matrix (PSSM) for CDK1 were taken from supplementary table 2 of ref. ^55^. These values are scaled such that a value of 1 represents an equal preference for all amino acids, >1 indicate a preference for an amino acid and <1 indicate negative selection against an amino acid. The serine(1.2371)/threonine(0.762) preference of the phospho-acceptor (position 0) was also added. To calculate the ICAS for a phosphosite, the preference scores for each residue in the −5 to +4 amino acid window were multiplied. For the analysis of human *in vitro* CKS-dependent phosphorylation, we combined the CKS-dependent cluster and CKS-dependent at high concentration cluster and compared them to the CKS independent cluster, as identified by the authors of that study^41^.

GO analyses were performed using ShinyGO^81^ (version 0.82) and PANTHER^82^. Enrichments were calculated based on hypergeometric distribution followed by false discovery rate (FDR) correction. Sequence Logos and Filled Sequence Logos were generated using the iceLogo server^83^. Substrate localisation was determined as described previously^84^.

CKS priming site annotation was performed using custom Python scripts, based on the reported criteria^37,38^. Phosphosites were annotated based on the presence or absence of a CKS binding motif ([FILPVWY]-X-T-P, 12-52 amino acids N-terminal, or T-P, 12-32 amino acids N-terminal) which was detected as phosphorylated in the same dataset. Sites were also annotated based on the presence of these motifs at equivalent distances in the C-terminal direction, or with serine as the phosphorylated residue, to analyse the directionality and specificity of possible priming mechanisms.

The timing of cell cycle phosphorylation for some substrates, as measured by the area under the curve of the phosphorylation profile over time (AUC), has been reported previously^4^.

### Data representation and statistical tests

Statistical tests were carried out using GraphPad Prism (v 10.1.1) and R (v4.3.1 and earlier). Distributions were compared using Mann-Whitney U tests where *n* = 2, and Kruskal-Wallis tests with post hoc Dunn’s test where *n* > 2. Paired data were compared using a paired T-test. Enrichment amongst clusters was determined using Fisher’s exact tests with Bonferroni correction for multiple testing. All tests were two-tailed. Where error bars are not visible they are masked by the data points.

## Data and code availability

All mass spectrometry data generated have been deposited to the ProteomeXchange Consortium via the PRIDE partner repository with dataset identifiers XXXX. Previously described custom scripts used to analyse fluorescent time-lapse data can be found at https://github.com/nkapadia27/SpatiotemporalOrchestration-of-Mitosis64.

## Acknowledgements

We thank A. Jones, H. Flynn and S. Howell for assistance with proteomics; T. Zeisner for the *mad2Δ synCut3-mCh* strain; S. Moreno for help with protein extraction protocols; J. Greenwood, T. Hammond, N. Kapadia, H. Kynman, L. Rider and T. Zeisner for critical reading of the manuscript; J. Hayles and all members of the Nurse lab for helpful discussions; and C. Gold for assistance when writing the manuscript. This work was supported by the Francis Crick Institute, which receives its core funding from Cancer Research UK (CC2003), the UK Medical Research Council (CC2003), and the Wellcome Trust (CC2003). In addition, this work was supported by a Wellcome Trust Investigator Award to P.N. (grant number 214183), The Lord Leonard and Lady Estelle Wolfson Foundation, and Woosnam Foundation.

## Author contributions

J.F.C. and P.N. initiated the study. J.F.C. designed and performed all experiments. S.B. generated the Suc1^ts^ strain. J.F.C. and T.A. performed and analysed IP mass spectrometry experiments. J.F.C. performed all other data analysis. J.F.C. and P.N. wrote the manuscript with input from all authors.

---

**Extended Data Fig. 1:**
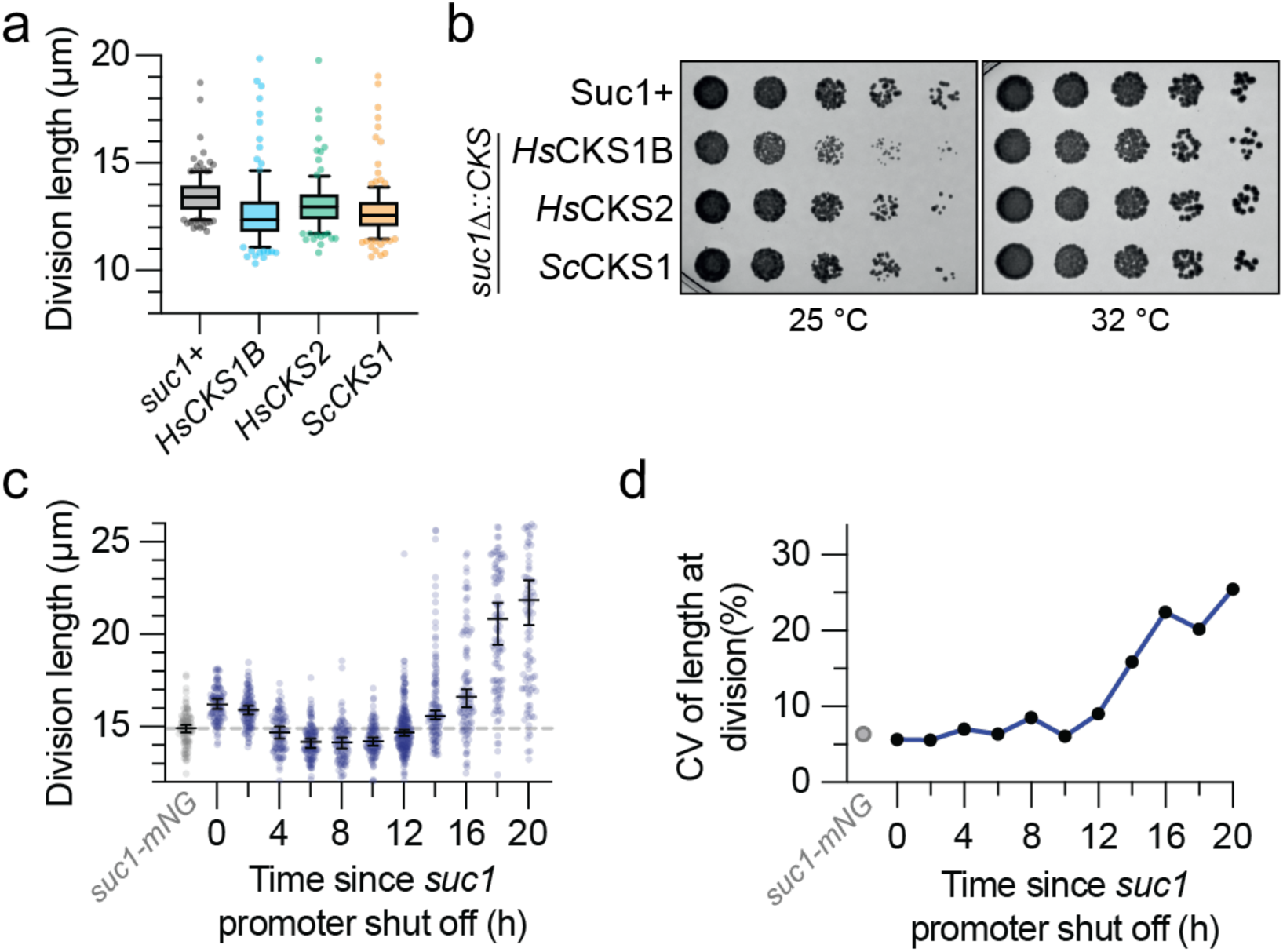
CKS protein function is conserved across eukaryotes. **a**, Length at septation for *suc1Δ::CKS* homolog strains. 4 data points are beyond the y axis upper limit. *n* > 100 cells per strain, *n* = 3 biological repeats. **b**, Representative serial dilution assay for colony formation of *suc1Δ::CKS* homolog strains. **c**, Length at septation and **d**, coefficient of variation in septation length following *suc1* promoter shut off. *n* > 100 cells per strain, *n* = 2 biological repeats. 55 data points are beyond the y axis upper limit.

**Extended Data Fig. 2:**
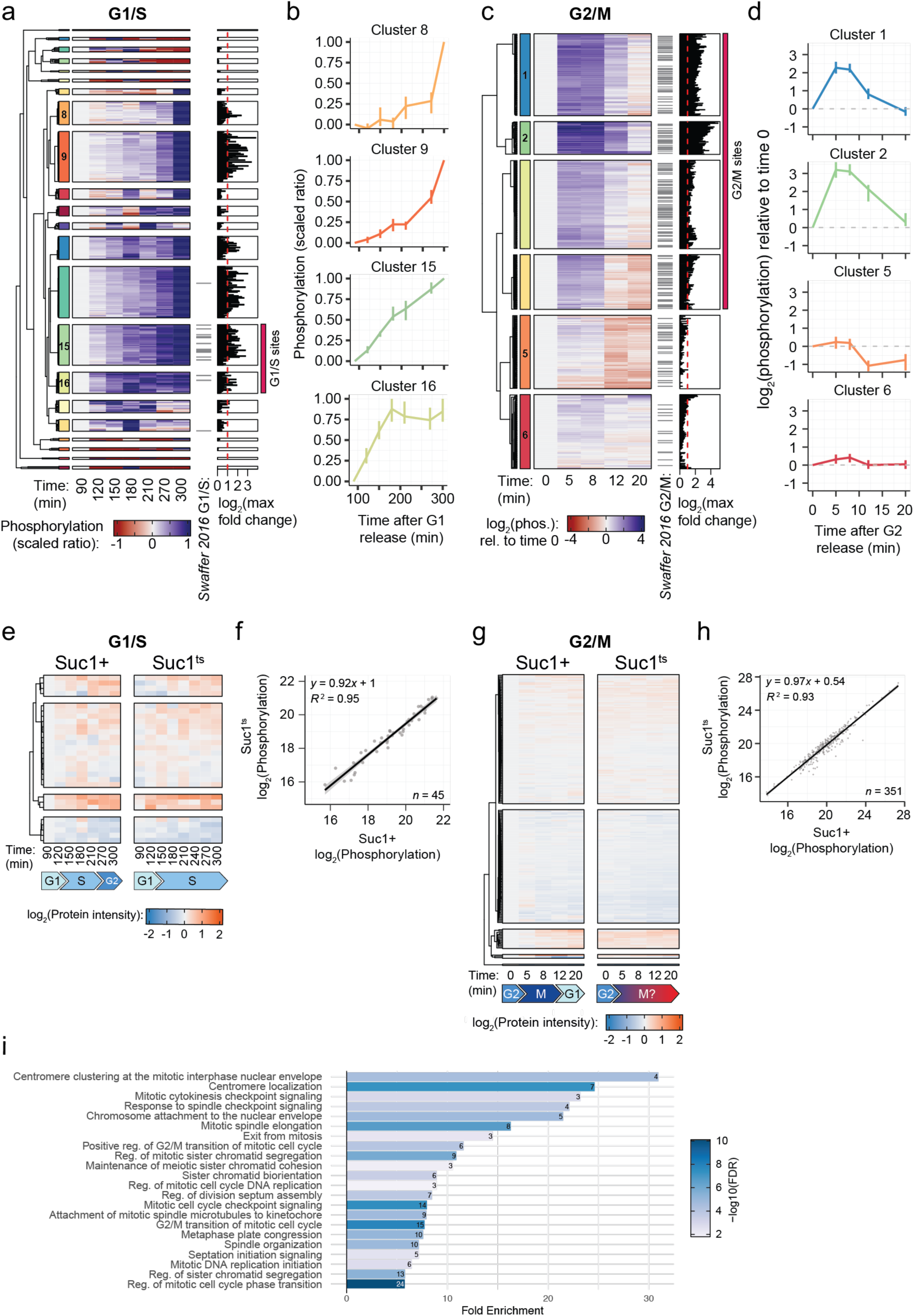
Identification of CDK phosphosites of interest at G1/S and G2/M. **a**, Heatmap of the phosphorylation intensity of 306 phosphosites which have been previously identified as CDK dependent (ref. ^4^ and unpublished) during progression through S-phase in the *suc1+* strain. Grey lines indicate phosphosites previously annotated as G1/S CDK substrates^4^. Maximum fold change in phosphorylation intensity across all time points is depicted as a black bar chart for each line in the heatmap. Sites were clustered by Euclidean distance. Red box indicates sites identified for further study as G1/S CDK phosphosites. **b**, Average phosphorylation behaviour (median and IQR) of selected clusters identified in **a**. Clusters 8 and 9 contain example mitotic CDK sites phosphorylated late in the time course, whereas Clusters 15 and 16 are phosphorylated during the G1/S transition. **c**, as in **a**, but for 569 phosphosites detected during progression through mitosis. **d**, Average phosphorylation behaviour (median and IQR) of selected clusters identified in **c**. Clusters 5 and 6 contain example early CDK sites which change little during mitosis, whereas Clusters 1 and 2 are phosphorylated during the G2/M transition. **e**, Heatmap of changes in protein intensities relative to time 90 min for the 28 proteins detected which contain G1/S CDK phosphosites of interest. Sites were clustered by Euclidean distance. **f**, Phosphorylation intensity in Suc1+ and Suc1^ts^ strains at time 90 min for the G1/S experiment. Black line and grey shading represent a least-squares linear regression fit with 95 % CIs. **g**, Heatmap of changes in protein intensities relative to time 0 min for the 205 proteins detected which contain G2/M CDK phosphosites of interest. Sites were clustered by Euclidean distance. **h** as in **f**, but comparing phosphorylation at time 0 min in the G2/M experiment. **i**, GO Biological Process enrichment for combined Suc1-dependent sites from G1/S and G2/M. Numbers inside bars indicate the number of genes in each GO annotation group. FDR – False Discovery Rate.

**Extended Data Fig. 3:**
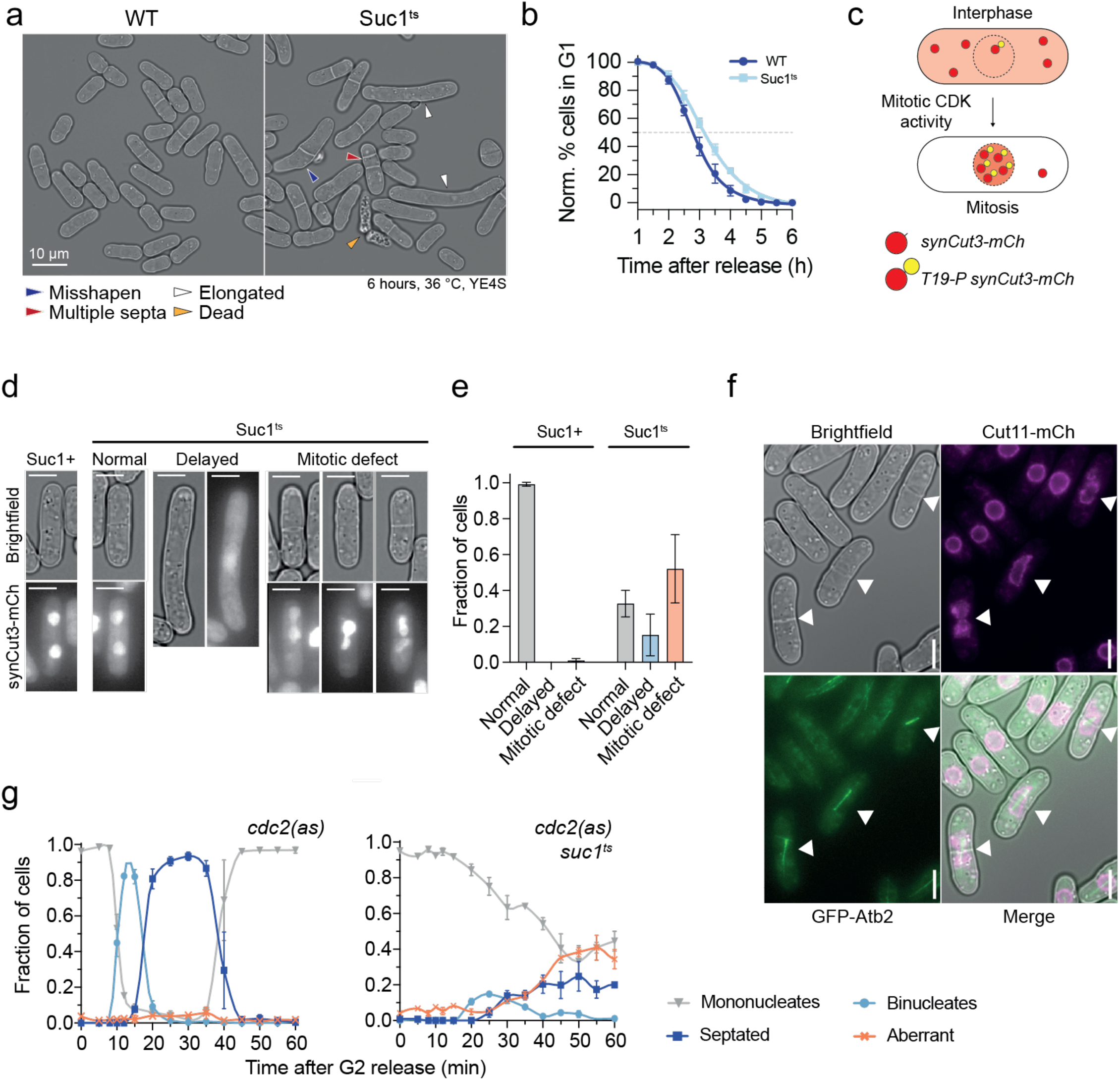
Suc1 regulates the onset and progression of both S-phase and mitosis. **a**, Example brightfield images of cells 6 h after shift to the restrictive temperature. Coloured arrows indicate assorted cell cycle defects. **b**, Quantification of data in Fig. 3a, indicating the normalised percentage of cells with 1C DNA content over time (Methods). Least-squares regression fit of a sigmoidal function. Mean and s.d. shown for *n* = 5 biological repeats. **c**, Schematic representation of the synCut3-mCh CDK activity sensor. **d**, Example brightfield and synCut3-mCh images illustrating the classes of behaviour observed. **e**, Quantification of the frequencies of the classes in **d.** Mean and range shown for *n* = 2 biological repeats. **f**, Example brightfield, alpha tubulin (GFP-Atb2), nuclear marker (Cut11-mCh) and merged channels. White triangles indicate short mitotic spindles in ‘cut’ cells, with division septa bisecting undivided nuclei. Scale bar = 5 µm. **g**, Binucleation and septation indices following release at the restrictive temperature from a G2 arrest. *n* > 100 cells per timepoint, Mean and range shown for *n* = 2 biological repeats. Lines represent Akima spline fits with 72 segments per strain.

**Extended Data Fig. 4:**
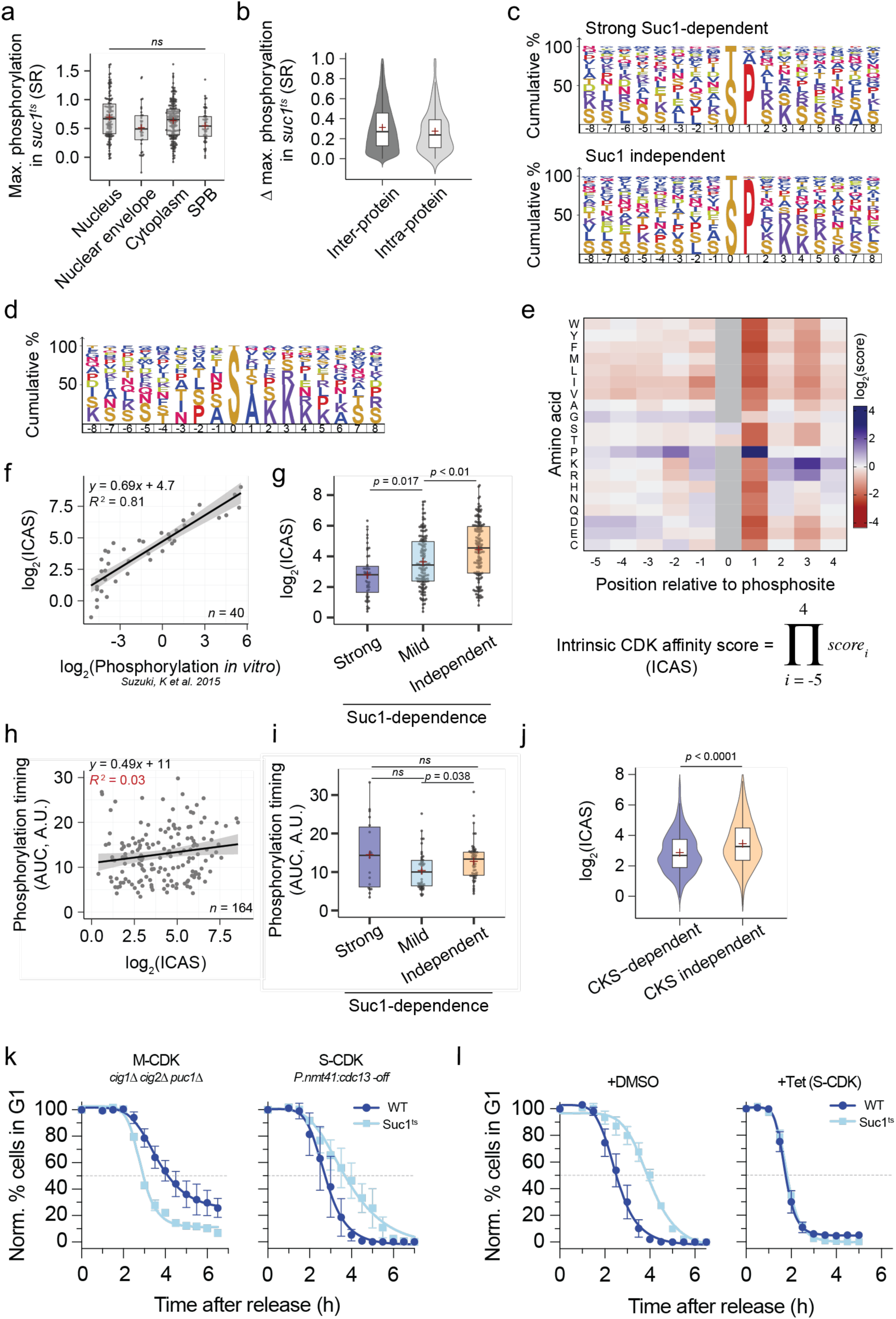
CKS enhances phosphorylation of intrinsically low-affinity phosphosites. **a**, Maximum phosphorylation reached in the Suc1^ts^ condition, grouped by substrate localisation (Methods). SR = Scaled ratio. 3 datapoints lie beyond the axis limits. *n* = 130, 35, 184, 51 for nucleus, nuclear envelope, cytoplasm and SPB (Centrosome) respectively. **b**, Distribution of the difference between the maximum phosphorylation reached in the Suc1^ts^ condition for phosphosite pairs on different proteins (inter-protein, *n* = 19,615) and phosphosite pairs on the same protein (intra-protein, *n* = 285). Difference values were only calculated for phosphosites from substrates with more than one phosphosite. Box plots represent IQR, with a black line at the median and a red + at the mean. Distributions are statistically different (p < 0.01) due to high test power, but means differ by only 0.05 (SR). (363 Inter-, and 2 Intra-data points are beyond the axes limits) **c**, Filled sequence logo representation of amino acid frequencies surrounding phosphosites for Strong Suc1-dependent and Suc1 independent phosphosites compiled from both phosphoproteomic experiments. **d**, Filled sequence logo representations of amino acid frequencies surrounding non-canonical phosphosites. **e**, Heatmap representing the “Site Score” for each amino acid at positions −5 to +4 surrounding a phosphosite for human CDK1^55^, and the equation for calculating Intrinsic CDK Affinity Score (ICAS). **f**, ICAS compared to *in vitro* phosphorylation intensity of peptides by *Xenopus* CDK1^52^. Least-squares fit of a linear regression. **g**, ICAS for canonical S/T-P G2/M phosphosites grouped by Suc1-dependence. Box plots represent IQR, with a black line at the median and a red + at the mean. *n* = 47, 133 and 137 for Strong, Mild and Suc1-independent respectively. **h**, Correlation between the timing of phosphorylation during the cell cycle, as determined by integration of phosphorylation intensity over time (AUC)^4^ and ICAS. Larger AUC indicates earlier cell cycle phosphorylation. Least-squares fit of a linear regression. **i**, Timing of phosphorylation during the cell cycle, as in **h**, grouped by Suc1-dependence. *n* = 43, 18 and 55 for Strong, Mild and Suc1-independent respectively. **j**, ICAS for *in vitro* phosphorylation data for canonical S/T-P phosphosites from ^41^ grouped by CKS-dependence, as determined in that study. Box plots represent IQR, with a black line at the median and a red + at the mean. *n* = 448, 1164 for CKS-dependent and independent groups, respectively. 4 data points lie outside the axes bounds. **k**, Normalised percentage of cells with 1C DNA content over time after release from a G1 arrest (Methods). S-phase was driven by the indicated Cyclin-CDK complexes, as in Fig. 4g. Least-squares regression fit of a sigmoidal function. Mean and range shown for *n* = 2 biological repeats. **l**, Normalised percentage of cells with 1C DNA content over time after release from a G1 arrest in the presence of just S-CDK activity (*P.nmt41:cdc13* + thiamine). Least-squares regression fit of a sigmoidal function. Tetracycline (Tet) addition results in over-expression of a Cig2-L-Cdc2 S-CDK fusion protein from a tetracycline-inducible promoter. Mean and s.d. shown for *n* = 3 biological repeats.

**Extended Data Fig. 5:**
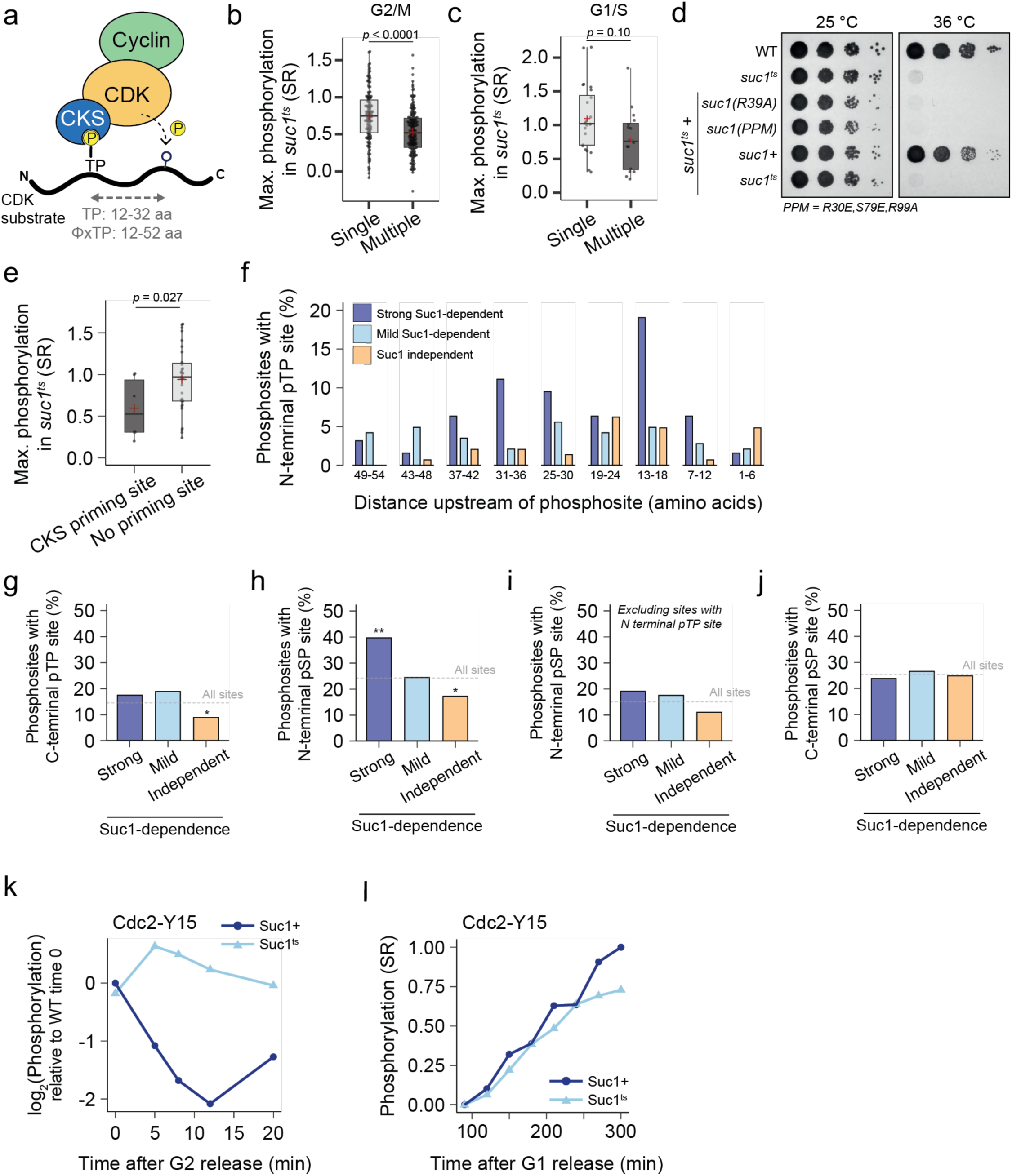
Suc1 enhances local and global CDK activity. **a**, Schematic representation of CKS-mediated priming mechanism by binding to N-terminal pT-P residues. ϕ - hydrophobic residue. **b** and **c**, Maximum phosphorylation reached in the Suc1^ts^ condition at G2/M (**b**) and G1/S (**c**), grouped by how many CDK sites have been identified on the protein substrate (Single, vs Multiple). Box plots represent IQR, with a black line at the median and a red + at the mean. SR = Scaled ratio. 2 datapoints lie beyond the axis limits in each panel. *n* = 151, 200 for single and multiple (**b**), and *n* = 25, 17 for single and multiple (**c**) respectively. **d**, Representative serial dilution assay for colony formation to assay if mutant *suc1* alleles can rescue the lethality of *suc1^ts^*. Except for the top two controls, strains are *suc1^ts^* with a second *suc1* allele *in trans*. R39A and PPM(R30E,S79E,R99A) are the analogous mutations to those from previously reported phospho-binding pocket mutants, based on protein structure^32,37^. **e**, Maximum phosphorylation reached in the Suc1^ts^ condition at G1/S, grouped by the presence or absence of a putative upstream CKS priming phosphosite. Box plots represent IQR, with a black line at the median and a red + at the mean. SR = Scaled ratio. 5 datapoints lie beyond the axis limits. *n* = 6, 36 for CKS priming and no priming respectively. **f**, Percentage of phosphosites at G2/M in each behaviour class with a putative upstream pT-P CKS-priming phosphosite, binned by distance between the phosphosite and putative priming site. **g**, Percentage of phosphosites at G2/M in each behaviour class with C-terminal pT-P phosphosites. Dashed line indicated the frequency across the entire dataset. **h**, as in **g**, but assessing N-terminal pS-P phosphosites, which are not predicted to bind CKS. **i**, as in **h**, but having eliminated sites with both an N-terminal pT-P and pS-P putative priming site. The observed enrichment collapses indicating that it is likely due to nearby pT-P sites and not direct pS-P priming by CKS. **j**, as in **g**, but assessing C-terminal pS-P phosphosites. **k**, and **l**, Phosphorylation of Cdc2-Y15 as detected by mass spectrometry from the G2/M (**k**) and G1/S (**l**) datasets. SR = Scaled Ratio.

**Supplementary Figure 1:**
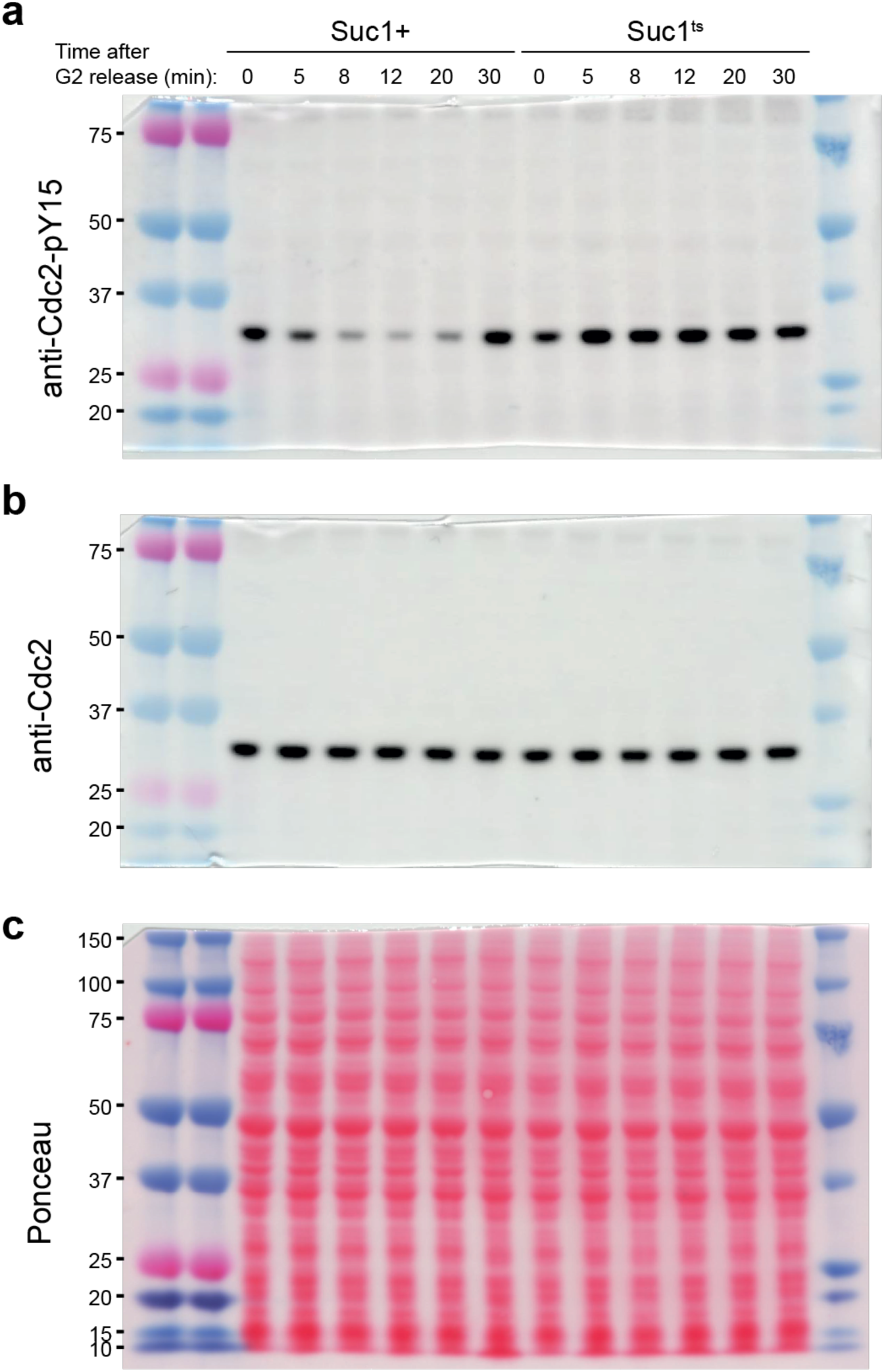
Uncropped western blots relating to Figure 5f. The lane annotations inpanel (a) apply to all panels. a) Western blot scan for Figure 5f upper panel. b) Western blot scan for Figure 5f middle panel. c) Ponceau-S strain scan for Figure 5f lower panel.

